# The novel conserved NAD^+^-binding micropeptide SGHRT regulates mitochondrial function and metabolism in human cardiomyocytes

**DOI:** 10.1101/2021.10.24.465637

**Authors:** Vinh Dang Do, Nikhil Kumar Tulsian, Warren KY Tan, Zhe Li, Liyi Cheng, Matias I. Autio, Wilson LW Tan, Zenia Tiang, Arnaud Perrin, Jianhong Ching, Mayin Lee, Isabelle Bonne, Chrishan Ramachandra, Choon Kiat Lim, Derek J Hausenloy, Chester Lee Drum, A. Mark Richards, Ganesh S. Anand, Roger SY Foo

**Affiliations:** Cardiovascular Research Institute, Yong Loo Lin School of Medicine, National University of Singapore (NUS); Cardiovascular Disease Translational Research Programme, National University Health Systems, Singapore; Genome Institute of Singapore, Agency of Science Research and Technology, Singapore; Department of Biological Sciences, Faculty of Science, National University of Singapore; Department of Biochemistry, Yong Loo Lin School of Medicine, National University of Singapore; Department of Cardiology, National Heart Centre Singapore, Singapore; Cardiovascular and Metabolic Disorders Program, Duke-NUS Medical School, Singapore; Life Sciences Institute Immunology Programme, Centre for Life Sciences, National University of Singapore; Cardiovascular ACP, Duke-NUS Medical School, Singapore; Christchurch Heart Institute, University of Otago, New Zealand; Department of Chemistry, PennState University, Pennsylvania, USA

## Abstract

Nicotinamide adenine dinucleotide (NAD) is a critical metabolite and coenzyme for multiple metabolic pathways and cellular processes (*1-4*). In this study, we identified Singheart, SGHRT as a nuclear genome-encoded NAD+-binding mitochondrial micropeptide. SGHRT, present in both monomeric and dimeric forms, binds directly to NAD, but not NADH or flavin adenine dinucleotide (FAD). Localized to the inner mitochondrial membrane and mitochondrial matrix, SGHRT interacts with the mitochondrial enzymes Succinate-CoA Ligase and Succinate Dehydrogenase. *SGHRT* deletion in human embryonic stem cell derived cardiomyocytes disrupted mitochondria morphology, decreased total NAD and ATP abundance, and resulted in defective TCA cycle metabolism, the electron transport chain and in Ox-Phos processes. These results comprise the first report of an NAD^+^-binding micropeptide, SGHRT, required for mitochondrial function and metabolism.

## Main Text

Emerging discoveries implicate multiple nuclear genome encoded micropeptides in diverse biological processes, including embryogenesis (*5*), myogenesis (*6, 7*), development (*8*), cellular metabolism (*9-13*), inflammation (*14, 15*), apoptosis (*16*), calcium handling (*17-19*) carcinogenesis (*20-22*), and multiple settings of disease pathogenesis (*19, 23, 24*). A large number of these micropeptides have been particularly identified in heart and muscle cells (*9-11, 17, 18, 25*), as small ORFs embedded in genes annotated as long noncoding RNA. Micropeptides are localized in different subcellular compartments including the nucleus (*26-30*), cytoplasm (*20, 21, 31*), endosome (*21, 32, 33*), SR (*6, 18, 34*), cell membrane (*7, 35*), and mitochondria (*9-13, 15, 16, 25, 36*), or may even be secreted (*5, 37*). Indeed, micropeptides that are largely enriched in mitochondria (*11, 25*) participate in fatty acid oxidation (*9, 10*), oxidative stress (*16*), metabolic homeostasis (*12, 36*), oxidative metabolism (*11, 25*), and in the unfolded protein response (UPR) (*38*). Our work provides the first report of a mitochondrial-localised micropeptide that binds to nicotinamide adenine dinucleotide (NAD+).

The novel long intergenic non-coding RNA, *Singheart* (*Sghrt*), was identified through single nuclear transcriptomes of mouse and human cardiomyocytes, and highly co-expressed in a genetic network of key cardiac genes (*39*). The human *SGHRT* locus harbours a small ORF encoding a 47-residue micropeptide previously named as short transmembrane mitochondrial protein 1 precursor, STMP1. A peptide sequence with 100% homology was isolated previously in a proteome screen of bovine cardiac mitochondrial fractions, co-purified with subunit 9 of mitochondrial respiratory complex III (*40*). Moreover, multiple sequence alignment demonstrates that the STMP1 micropeptide sequence is highly conserved across vertebrates (*41*). Morpholino-based targeting in zebrafish produced a phenotype that included cardiac edema (*41*). More recently, it was also reported that the mouse *Sghrt* genomic locus encodes a novel mitochondrial micropeptide, namely Mm47, required for the activation of the NLRP3 inflammasome in macrophages (*14*). We therefore set out to characterize cardiac SGHRT micropeptide, and elucidate its role in mitochondrial function using human cardiomyocytes.

### SGHRT is a novel conserved mitochondrial micropeptide that exists in monomers and dimers

To confirm and resolve the protein expression of cardiac SGHRT, we performed mass spectrometry analysis with mouse, rat, pig and human hearts. Samples were prepared using a gel-assisted sample preparation (GASP) strategy (*42*), and peptide mapping was carried out with mitochondria enriched heart lysates and human embryonic stem cell (hES)-derived cardiomyocytes. To increase protein coverage, LC-MS/MS was performed in undigested, trypsin digested, and chymotrypsin digested samples. SGHRT peptides were confirmed, with full coverage achieved in rat and mouse samples (Fig. 1A, B). The predicted size of SGHRT micropeptide is ∼5-6 KDa, consistent with previous studies (*14*). However, when SGHRT synthesized peptides were resolved in 16.5% Tricine SDS gel, three bands were observed at ∼5.8, 8 and 12 kDa, with the latter band being most prominent. This result suggested that SGHRT oligomerises into dimers. The SGHRT oligomer was also resistant to heat (Fig. S1A) and EDTA treatment (Fig. S1B), suggesting a highly stable complex. The molecular weight of SGHRT peptides was further analysed quantitatively by analytical ultracentrifugation (AUC), which showed peaks in the range of 15-20 KDa, with the most abundant peak at ∼17 kDa (Fig. S1C). SGHRT oligomerisation was further validated *in vivo* by co-immunoprecipitation using human SGHRT that was C-terminally tagged with either FLAG or HA epitopes, transfected in HEK cells (Fig. 1C, D). In addition, co-staining with antibodies against FLAG and HA confirmed co-localization of SGHRT-FLAG and SGHRT-HA (Fig. 1E). Next, using alanine scanning mutagenesis (Fig. S1D-E), we identified that the glycine residues (G) at positions 7, 11 and 15 are important residues for dimerization, and form a zipper motif GxxxGxxxG (*43*). Notably, asparagine (N21) that stabilizes isopeptide bonds with adjacent lysine (L) in certain bacterial cell-surface proteins (*44*), and proline (P25) mutants also resulted in loss of the dimer band. These residues are clearly important for SGHRT dimerization (Fig. 1F) Together, these data indicate that SGHRT encodes a conserved mitochondrial micropeptide that exists in both dimeric and monomeric forms.

**Fig. 1.**
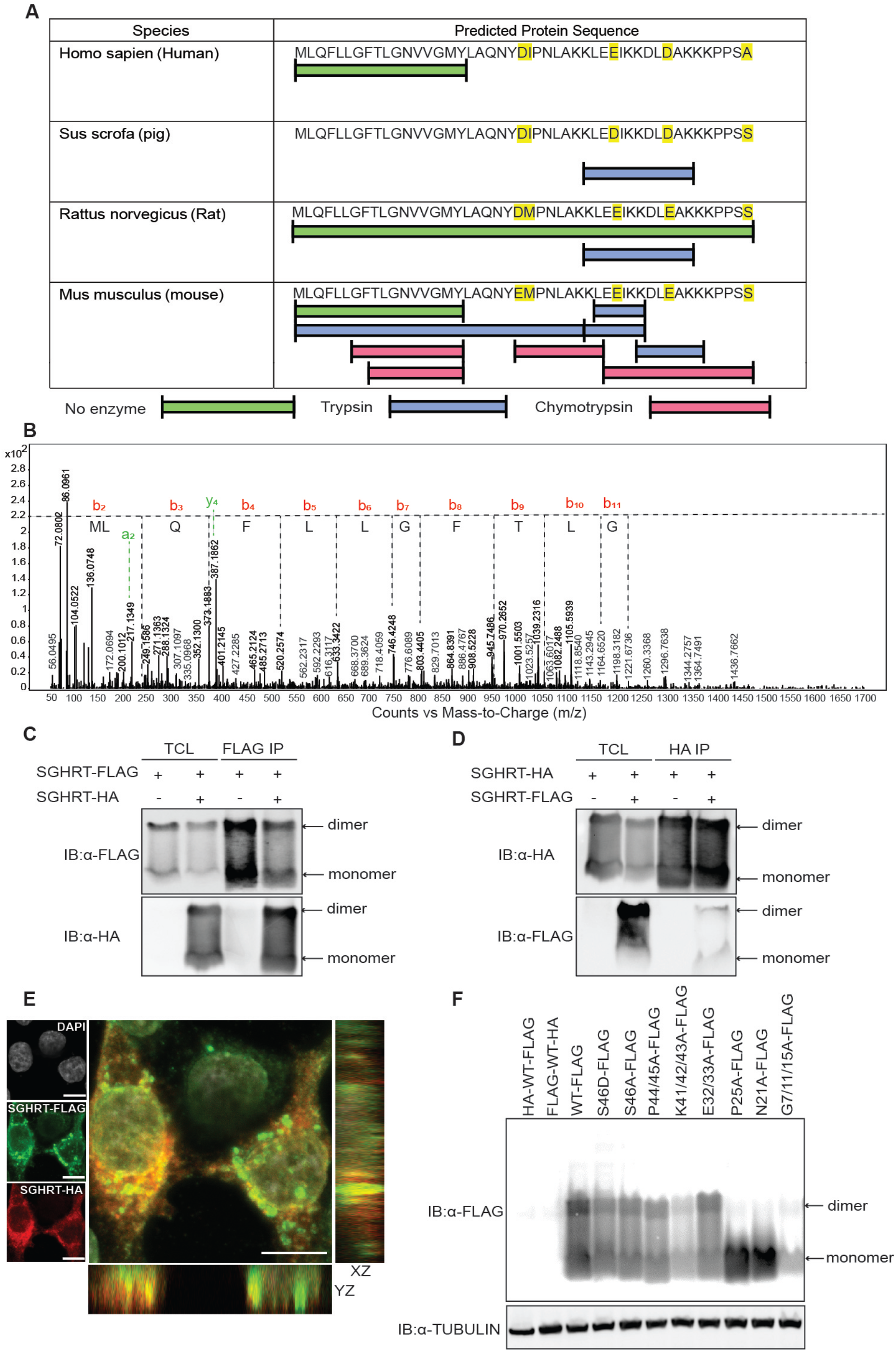
Identification of SGHRT micropeptides that form oligomers. (**A**) LC-MS/MS peptide mapping of SGHRT using undigested, trypsin digested, and chymotrypsin digested mammalian mitochondrial preparations. Residues highlighted in yellow are species specific. Full length peptide was detected in undigested rat mitochondrial preparations. (**B**) Representative MS/MS spectrum of intact SGHRT in rat preparations. (**C, D**) Total cell lysate (TCL) showed FLAG- and HA-tagged SGHRT displayed as monomers and dimers on Western blot when expressed in HEK293 cells. Co-immunoprecipitation confirmed interaction between FLAG- and HA-tagged SGHRT as oligomers. (**E**) Immunofluorescence with anti-FLAG (green) and anti-HA (red) antibodies in transfected HEK293 cells validated cytoplasmic cellular co-localisation of SGHRT peptides. Scale bar, 10 µm. (**F**) Western blot of transfected alanine-mutant and WT SGHRT constructs in HEK293 cells showed residues Pro25, Asp21 and Gly7/11/15 to be necessary for oligomerisation. Anti-tubulin antibody was used as loading control.

### SGHRT binds directly to NAD^+^ but not FAD or NADH

The GxxxG motif in the SGHRT N-terminus (Fig. S1D) has been described before as an adenine dinucleotide binding motif (6), and its homologs were shown to localized to the mitochondrial membrane. We therefore tested the likelihood of SGHRT binding to flavin adenine dinucleotide (FAD), nicotinamide adenine dinucleotide (NAD^+^) or its reduced form (NADH). First, we monitored the conformational dynamics of full-length SGHRT synthetic peptide alone and complemented it with the intrinsic dynamics of smaller fragments. Amide Hydrogen-Deuterium Exchange Mass Spectrometry (HDXMS) of full-length SGHRT synthetic peptide revealed a global uptake of 33% and 43% at 1 min and 10 min of deuterium labeling, respectively (Fig. 2A-E). This corroborated with average relative fractional uptake (35%) of pepsin-proteolyzed fragments of SGHRT (Fig. S2A-B). Next, we performed HDXMS analysis of SGHRT peptide saturated with FAD, NAD^+^ or NADH. NAD^+^ binding resulted in large-scale decreases in global deuterium exchange of full-length SGHRT (Fig. 2C), but not with FAD (Fig. 2B) nor NADH (Fig. 2D), indicating SGHRT binds specifically to NAD^+^, and not NADH or FAD.

**Fig. 2.**
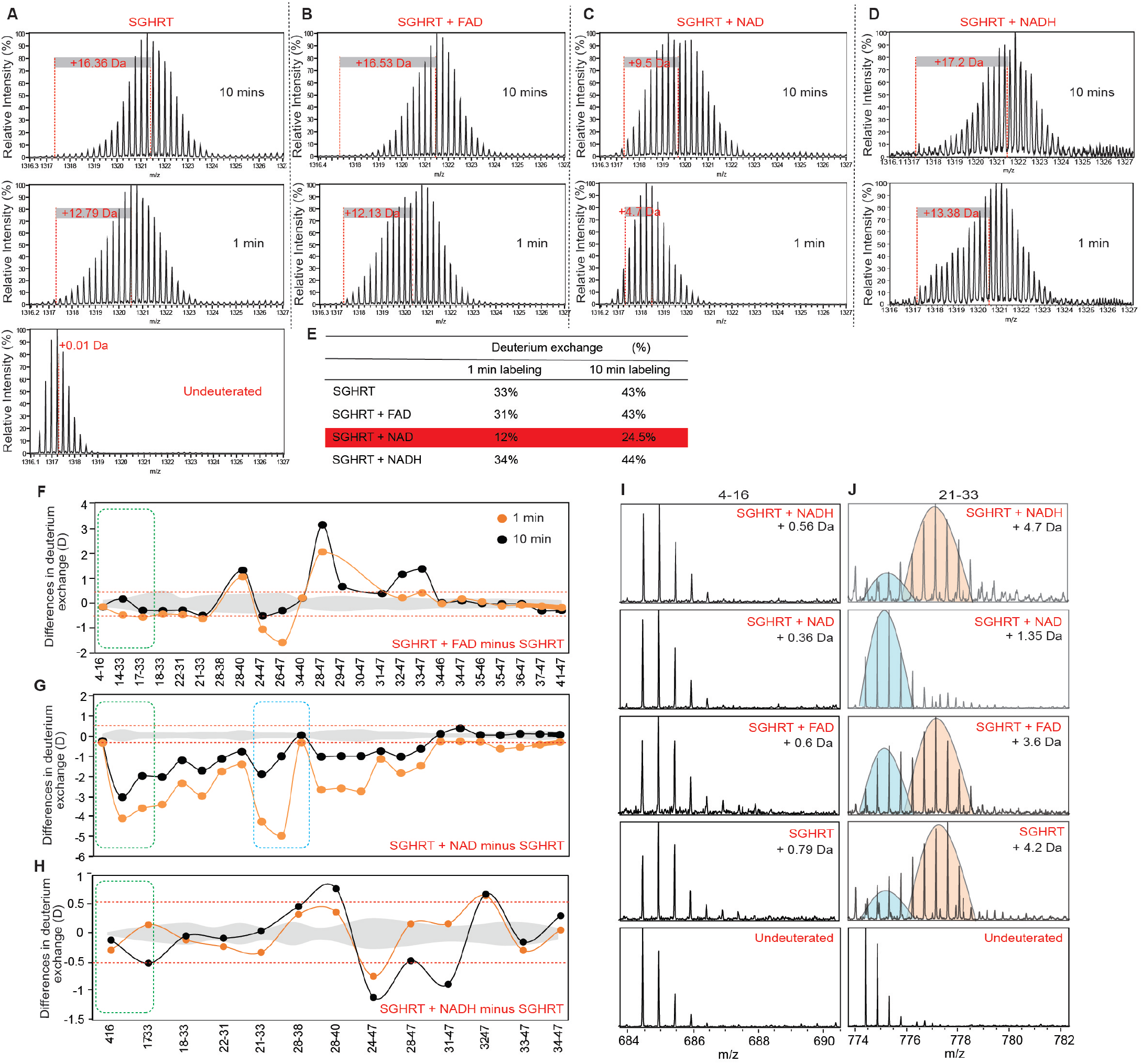
HDX-MS of pepsin-digested and undigested SGHRT micropeptides in the presence of FAD, NAD and NADH. (**A-D**) Mass spectral plots showing the global HDX-MS readout of SGHRT micropeptide alone (**A**), in the presence of FAD (**B**), NAD (**C**) and NADH (**D**) at 1 min and 10 min of deuterium labeling. Deuterium exchange centroid values are calculated as described in materials and methods. Mass spectrum of undeuterated control is shown for reference. Incubation with NAD showed significant decrease in global deuterium exchange of SGHRT (**C**). (**E**) Table summarizing the percentage of deuterium exchange at 1 and 10 min labelling times. (**F-H**) HDXMS readouts of SGHRT for different pepsin-proteolyzed fragments indicated by their residue numbers (X axis). Differences in deuterium exchange (Y-axis) of SGHRT, in the presence of FAD (**F**), NAD (**G**) or NADH (**H**), following subtraction of free SGHRT. Positive differences indicate increased deuterium exchange, and negative differences indicate decreased deuterium exchange in SGHRT bound to FAD, NAD or NADH, as compared to SGHRT alone. A significance threshold of (±0.5 D) is indicated by red-dotted line and standard deviations are in gray. Green-dashed and blue-dashed boxes highlight important amino acid residues that reflect binding to NAD, and not FAD or NADH. (**I, J**) Stacked mass spectral plots for SGHRT, SGHRT+FAD, SGHRT+NAD, and SGHRT+NADH at 1 min labeling time for pepsin-proteolyzed fragment specifically for amino acid residues spanning 4-16 (**I**) and 21-33 (**J**). In (**J**), the 21-33 residue SGHRT fragment showed characteristic bimodal spectra distribution (more of high deuteration: right-shifted peak in orange; small low deuteration peak in cyan), with and without FAD or NADH, but a significantly lower exchange and unimodal distribution (in cyan only) with NAD, indicating direct binding of the 21-33 fragment with NAD. Mass spectra of undeuterated experiment is shown for reference.

Subsequently, to identify the residues involved in binding, we used smaller fragments of SGHRT after pepsin proteolysis. HDXMS results of SGHRT with FAD (Fig. 2F) or NADH (Fig. 2H) showed no major changes in deuterium exchange. Consistent with the global-HDXMS results above, NAD^+^ binding resulted in a significant reduction in deuterium uptake in distinct segments of SGHRT (Fig. 2G). Pepsin-digest fragments spanning the GxxxG motif (Fig 2G-green box) showed marginal differences (−0.3 D) in deuterium exchange. Peptides spanning 14-33 and 38-47 showed large-scale decrease in deuterium exchange (Fig. 2G-green and blue boxes) in the presence of NAD+. These results validate that NAD^+^ binds and induces conformational rigidity of SGHRT. Importantly, deuteration level changes were more pronounced at shorter labeling time (1 min), and less prominent at longer labeling time (10 min), indicating that NAD^+^ and SGHRT do not bind in stable fashion for longer durations (Fig. 2G), which can be attributed to its function as a NAD+ carrier. Examining the isotopic distribution profiles of peptide spanning 21-33 (Fig. 2J) showed the contrasting effects of NAD+ on SGHRT, compared to FAD or NADH. Taken together, the findings indicate SGHRT binds specifically to NAD+ but not NADH or FAD, and SGHRT residues 14-33 are important for NAD^+^ binding.

### SGHRT interacts with Succinate-CoA Ligases (SUCL) and Succinate Dehydrogenases (SDH)

Prior data and peptide analysis predicted SGHRT to be localised to the mitochondria. We therefore generated a hES cell line overexpressing SGHRT-FLAG using lentiviral infection. In mitochondria isolated from SGHRT-FLAG-overexpressing cells, progressive proteolysis with proteinase K and serial dilutions of digitonin confirmed that SGHRT monomers localized to the Inner Mitochondrial Membrane (IMM) and mitochondrial matrix (Fig. S3L). Although SGHRT lacks a classical mitochondrial targeting sequence, it contains a conserved domain comprising of a hydrophobic core next to charged amino acid residues (Fig. S1D, E), shown to be sequence determinants of IMM proteins (*45*). Next, to identify SGHRT interacting proteins, we generated a transgenic HEK293T cell line overexpressing SGHRT-3x FLAG or 3x FLAG (control) using lentiviral infection. Using pulldown against mouse heart left ventricular lysates, SGHRT interacting proteins were semi-quantified by LC-MS/MS and peptide mass fingerprinting (Fig. 3A). Network and Gene Ontology (GO) analyses revealed interactions primarily with mitochondrial proteins (Fig. 3B-C, Table S1). Co-localisation of SGHRT-FLAG to IMM and matrix succinate-coA ligases (SUCLG1, SUCLA2 and SUCLG2), as well as succinate dehydrogenases (SDHB and SDHA) were confirmed by immunostaining in both hES cells and hES-derived cardiomyocytes (Fig. 3D-H, Fig. S3A-E). On the other hand, SGHRT did not co-localise with NAD^+^-dependent dehydrogenases, IDH2 (Fig. S3, F and G) and OGDH (Fig. S3, H and I), and the NAD^+^ transporter SLC25A1 (*46, 47*) (Fig. S3J, K). To test whether SGHRT dimerization is important for its protein interactions, HEK293 cells were co-transfected with HA-SUCLG1 plasmid together with wildtype SGHRT-FLAG or dimer-deficient mutants of SGHRT-FLAG (G7/11/15A-SGHRT-FLAG, N21A-SGHRT-FLAG and P25A-SGHRT-FLAG). Co-immunoprecipitation revealed that SGHRT interaction with SUCLG1 was not affected by dimerization mutation (Fig. 3I). These observations were further supported by co-staining data using anti-FLAG and anti-HA antibodies (Fig. 3J). Similar results were observed for SGHRT and SDHA (Fig. 3K-L). Together, these data show that IMM-localised SGHRT monomers are sufficient to interact with SUCL and SDH proteins. Notably, mutational analyses of NAD+ binding residues did not perturb SGHRT interactions with other proteins.

**Fig. 3:**
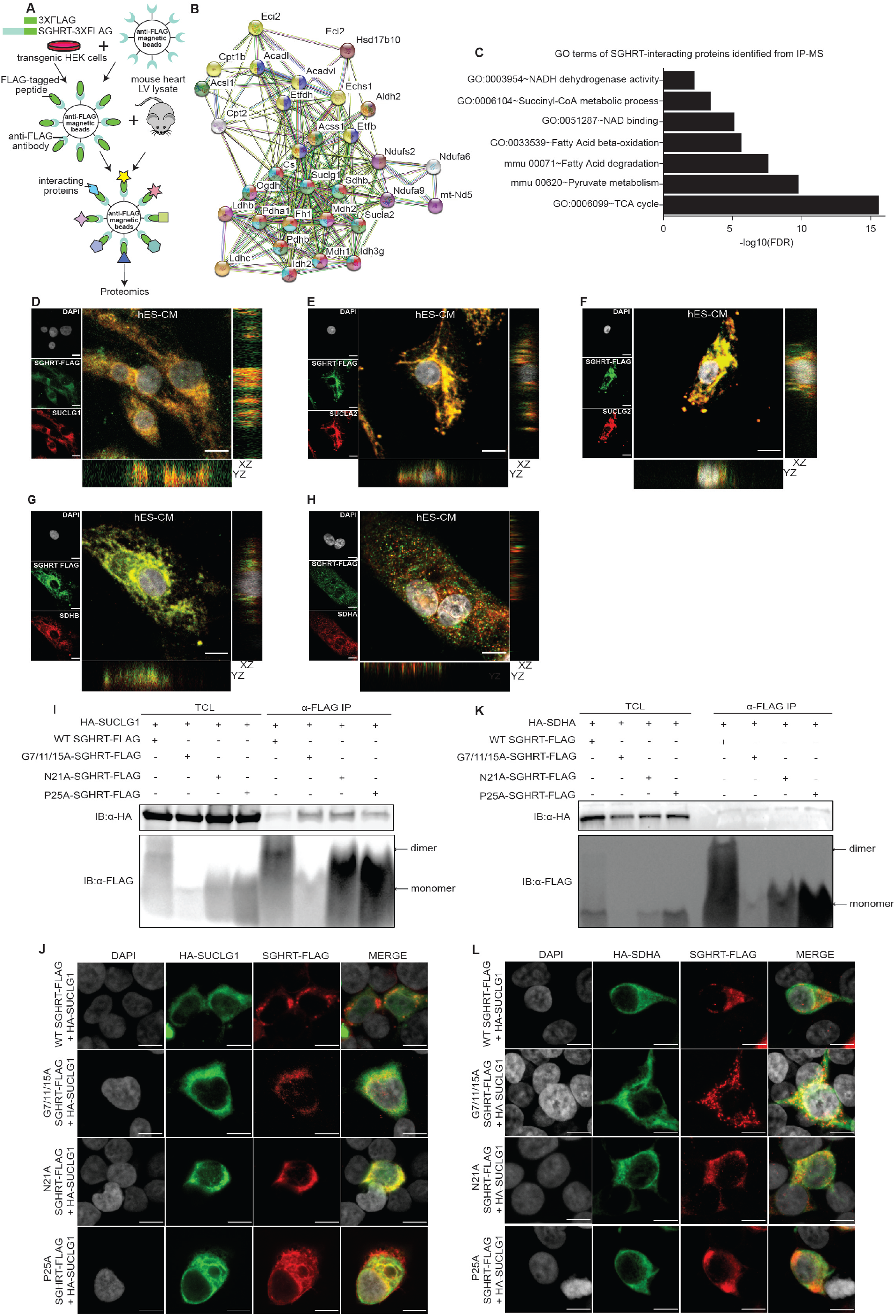
SGHRT interacts with mitochondrial proteins. (**A**) Schematic depicting the workflow used to identify SGHRT-interacting proteins. (**B-C**) STRING analysis (**B**) and GO analyses (**C**) of protein-protein interaction network indicating mitochondrial proteins are enriched in SGHRT-3XFLAG pulled down elute. (**D-H**) Immunofluorescence of SGHRT-FLAG (green) and TCA-related proteins of the mitochondria (red) including SUCLG1 (**D**), SUCLA2 (**E**), SUCLG2 (**F**), SDHB (**G**), SDHA (**H**), identified from IP/proteomics data in SGHRT-FLAG hES-CM. Scale bar, 10 µm. (**I, J**) HEK293 cells were co-transfected with HA-SUCLG1 plasmid together with wildtype SGHRT-FLAG or alanine mutated SGHRT-FLAG (G7/11/15A-SGHRT-FLAG, N21A-SGHRT-FLAG and P25A-SGHRT-FLAG) that disrupt dimerization. Immunoprecipitation was done using anti-FLAG, and western blot of total cell lysis and IP elute was performed with anti-FLAG and anti-HA antibodies (**I**), whereas Immunofluorescence was also done using anti-FLAG (green) and anti-HA (red) antibodies, scale bar, 10 µm (**J**). (**K, L**) HEK293 cells were cotransfected with HA-SDHA plasmid together with wildtype SGHRT-FLAG or alanine mutated SGHRT-FLAG (G7/11/15A-SGHRT-FLAG, N21A-SGHRT-FLAG and P25A-SGHRT-FLAG). Immunoprecipitation was done using anti-FLAG, and western blot of total cell lysis and IP elute was performed with anti-FLAG and anti-HA antibodies (**K**), whereas Immunofluorescence was done using anti-FLAG (green) and anti-HA (red) antibodies, scale bar, 10 µm (**L**).

### SGHRT is required for mitochondrial function and metabolism in human cardiomyocytes

To assess functional relevance of SGHRT, knockout (KO) hES cell lines were generated using CRISPR/CAS9 targeted deletion of the *SGHRT* promoter and first exon. Efficient genomic deletion was validated by a complete loss of *SGHRT* mRNA expression (Fig. S4H). By immunostaining and Western blot analysis, we verified that there was no change to the pluripotency OCT4 marker (Fig. S4A, B). Cardiac differentiation based on a published differentiation protocol (*48*) proceeded normally with wild-type (WT) and KO cells, but at weeks 4 and 8, *SGHRT* KO cardiomyocytes (tracked by the cardiomyocyte *MYH6*-GFP reporter) appeared rounder, smaller, and more dispersed in culture, compared to WT (Fig. 4A). GO analysis of differential transcriptomes showed that the genes involved in mitochondrial respiration, glycolysis and fatty acid oxidation (FAO) were significantly downregulated in KO cardiomyocytes, compared to WT (Fig. 4B, Table S2). In addition, proteomic analysis by iTRAQ reflected a global reduction in expression of proteins associated with mitochondrial, oxidoreductase, and respiratory chain complexes (Fig. 4C, Table S3). Three-dimensional engineered heart tissues (EHT) (*49*) constructed using WT and KO D16 and D27 hES-cardiomyocytes showed a reproducible loss in tissue integrity (Fig. 4D, Movies. S1-4) and grossly abnormal contractility (Fig. 4E). GO analysis of differential transcriptomes of D7- and D16-KO EHTs were again consistent with the significant downregulation of genes in the redox process, cellular glucose homeostasis, mitochondrial transport and FAO, compared to WT (Fig. 4H, Fig. S4E, Table S4-S5). Indeed, scanning electron microscopy indicated abnormal mitochondria, with loss of distinct cristae structure in KO cardiomyocytes (Fig. 4G).

**Fig. 4:**
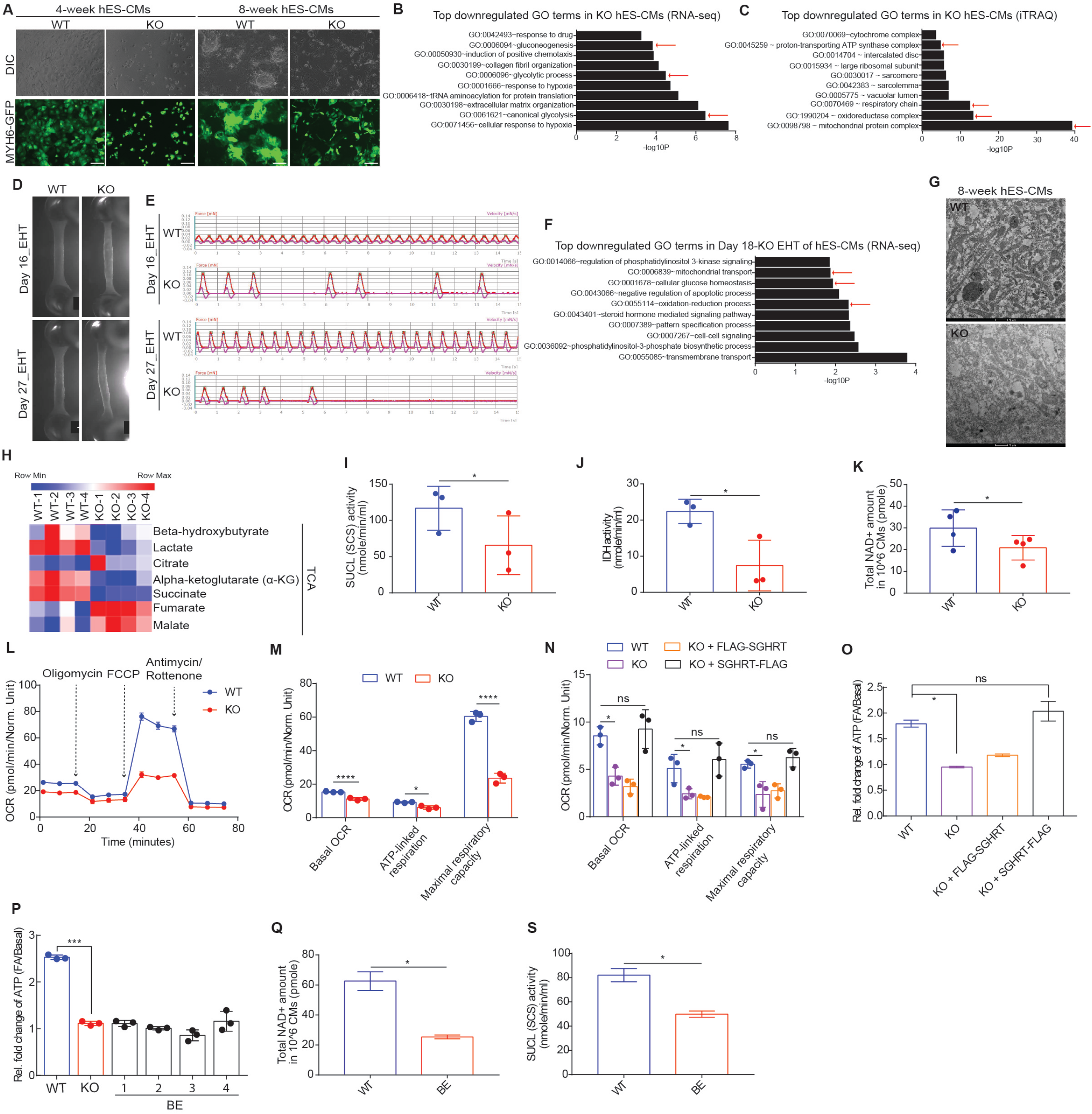
SGHRT is required for TCA metabolism and mitochondrial function in human cardiomyocytes. (**A**) Microscopic images showing morphological difference between WT and KO hES-CMs at 4-week and 8-week cardiac differentiation. (**B**) Gene ontology (GO) analysis of RNA-seq data showing top down-regulated of GO-term in 4-week KO hES-CM. (**C**) GO analysis of iTRAQ data showing top down-regulated GO terms in 4-week KO hES-CM. (**D, E**) Representative pictures showing *SGHRT* KO EHTs have poorly formed EHT profile (**D**) and abnormal contractility profile (**E**) compared to WT EHTs at D16 and D27 in culture. (**F**) GO analysis of RNA-seq data showing top down-regulated GO terms in day-16 KO EHTs. (**G**) Electron microscopic pictures of 8-week WT and KO hES-CMs, scale bar, 1 µm. (**H**) A heatmap showing expression of TCA metabolites in 4-week WT vs KO hES-CMs. (**I, J**) Graphs showing enzymatic activities of Succinyl-CoA Synthetase (SUCLA) (I), Isocitrate Dehydrogenase (IDH) (J) in 4-week WT and KO hES-CMs. (**K**) Graph showing total NAD level in 4-week WT and KO hES-CMs. (**L**) Seahorse Flux assay in 4-week WT and KO hES-CMs cultured in medium containing RPMI, B27 with insulin and 2 % FBS. (**M**) Quantitation of OCR for Seahorse in (L). (**N**) Quantitation of OCR of Seahorse Flux assay in 4-week WT, KO, KO with FLAG-SGHRT and KO with SGHRT-FLAG hES-CMs treated with 100µM fatty acid solution containing 100 μM of oleic acid and palmitic acid in a 1:1 ratio. (**O**) Quantification of ATP production in WT, KO, KO with FLAG-SGHRT and KO with SGHRT-FLAG with and without Fatty Acid treatment. (**P**) Quantification of ATP production in WT, KO and BE ES-derived CM clones with and without Fatty Acid treatment. (**Q, S**) Graphs showing total NAD level (**Q**) and enzymatic activity of SUCL (**S**) in 4-week WT and BE hES-CMs. Data are represented as mean ± SD from at least 2 biological replicates. *p < 0.05, **p < 0.01, **** p <0.0001, ns: not significant. Student’s t test with Bonferroni correction.

Further, targeted metabolomics showed significant reduction in KO cells for succinate and α-ketoglutarate (α-KG), with concurrent increased levels of malate and fumarate (Fig. 4H), reflecting an imbalance in tricarboxylic acid (TCA) cycle substrates and products, likely tilted by perturbed NAD^+^-dependent IDH enzyme activity. Direct *in vitro* enzymatic assays therefore confirmed defects in Isocitrate Dehydrogenase (IDH, Fig. 4I) and SUCL (Fig. 4J) function, but not of SDH (Fig. S4F), α-KG dehydrogenase (KGDH) (Fig. S4D) or Pyruvate Dehydrogenase (PDH) (Fig. 4SE). The reduction of IDH and SUCL enzymatic activities may explain reduced α-KG and succinate, which leads to the accumulation of fumarate and malate (Fig. 5SA, B). Although, why only one NAD^+^-dependent dehydrogenase is affected and not the others remains unclear. Indeed, total NAD (NAD^+^ and NADH) levels are also SGHRT-dependent (Fig. 4K), suggesting that SGHRT may be important for maintaining overall NAD homeostasis. Seahorse functional mitochondrial assay further confirmed that *SGHRT* KO impaired basal and maximal respiratory capacity in cardiomyocytes, cultured in both glucose-rich (Fig. 4L, M) or fatty acid-containing media (Fig. 4N). Similarly, *SGHRT* KO led to significant total ATP reduction (Fig. 4O), despite unchanged mitochondrial membrane potential (Fig. S4G). Importantly, the mitochondrial defects of *SGHRT* KO were completely rescued upon re-introduction of SGHRT-FLAG into KO cells (Fig. 4N-O, Fig. S4H).

To confirm that the phenotype in *SGHRT* KO resulted specifically from the loss of SGHRT micropeptide, we also generated ATG-mutated hES cell lines using CRISPR/CAS9 base editing of A or C to G or T, respectively (*50*). DNA sequencing analysis confirmed successfully two A-to-G (BE-1 and BE-2), and two C-to-T (BE-3 and BE-4) base-edited (BE) cell clones (Fig. S4I). While there was no significant change in *SGHRT* transcripts (Fig. S4J), SGHRT protein was lost in BE cells, compared to WT (Fig. S4K). Like *SGHRT* KO cells, ATP generation was blunted in BE cells (Fig. 4P). In addition, *SGHRT* BE cells also showed significant reduction in total NAD (Fig. 4Q) and SUCL enzymatic activity (Fig. 4S). Altogether, these results confirm that SGHRT micropeptide is responsible and necessary for mitochondrial function and metabolism, although the contribution of *SGHRT* transcript as a long noncoding RNA cannot yet be excluded.

In summary, we provide the first description of a conserved NAD^+^-binding micropeptide, SGHRT, localized to mictochondria and necessary for mitochondrial function and metabolism (Fig. S5). SGHRT micropeptides exist as both monomers and dimers, and its GxxxG motif, and residues N-21 and P-25 are important for dimerization. Dimerization appears to be unrelated to its IMM localisation, nor to its protein interactions with at least SUCL and SDH enzymes. SGHRT micropeptide binds directly to NAD+ via residues 14-33. Loss of SGHRT decreased total NAD+ levels in human cardiomyocytes, and reduced SUCL and IDH enzymatic activities.

Besides the importance of NAD+ in redox reactions, NAD+ is also an essential cofactor for non-redox NAD+ dependent enzymes, including SIRTUINs (*51*) and poly(ADP-ribose) polymerases (PARPs) (*52*). Further studies are warranted to identify the role of SGHRT in other mitochondria-localised NAD+ dependent functions of SIRTUIN-mediated deacetylation and PARylation. NAD+ is synthesized, catabolized and recycled in the cell constantly to maintain stable intracellular NAD+ levels in homeostasis. NAD synthesis pathways include *de novo* NAD synthesis from tryptophan, the salvage pathway via recycling or incorporation of nicotinamide, and the Preiss-Handler pathway using dietary nicotinic acid and the enzyme niconinic acid phosphoribosyltransferase (NAPRT), whereas NAD+ consumption is controlled by three main classes of NAD+-consuming enzymes including protein deacylases SIRTUINs, Poly(ADP-ribose) polymerases PARPs, and NAD+ glycohydrolases and cyclic ADP-ribose (cADPR) synthases (CD38, CD157 and SARM1) (*1, 2*). Interestingly, NAD+ metabolism is rapid and dynamic since its half-life varies between 15 mins to 15 hrs across different tissues (*53*). Our results showed that *SGHRT* deletion resulted in an overall reduction in total cellular NAD+, and a deeper understanding of this effect may provide insights into the molecular mechanisms of the ‘‘futile cycle’’ of NAD synthesis and degradation (*2*).

Finally, many micropeptides localized to the mitochondria interact with OXPHOS subunits (*9-11, 25*) and their mitochondrial localization of micropeptides may be facilitated by as yet undetermined localization signals or import systems, or may be purely driven by their small sizes and positively charged amino acid composition (*54*). Our results show that SGHRT is localized to mitochondrial IMM, interacting at least with SDH enzymes in complex II and SUCL enzymes, despite lacking obvious mitochondrial targeting peptide signals. Future studies may also reveal how SGHRT and other micropeptides are transported and localised to mitochondria. Nonetheless, our study reports for the first time a micropeptide SGHRT that critically regulates mitochondrial redox metabolism, including the TCA cycle, electron transport chain and OXPHOS processes, through direct binding with NAD+ in the mitochondria.

## Acknowledgment

We acknowledge Dr. Gao Bin and Dr. Ng Huck Hui for support with the *MYH6*-GFP hES cells, Dr. Liew Oi Wah for her support with the protein studies, Dr Jean-Paul Kovalik for discussions on mitochondrial metabolism, Dr. Zing Tan Tsze Yin for her support with IC-MS, and Dr. Siddharth Deshpande for his support with the AUC experiment.

## Author contributions

Conceptualization: RSYF, VDD. Methodology and experiments: VDD, NKT, WLWT, WKYT, ZL, LC, MIA, ZT, AP, JC, ML, IB, CR, CKL. Data and result analysis: VDD, RSYF, WLWT, DJH, CLD, MR, GSA. Manuscript writing: VDD, RSYF

## Data and materials availability

Plasmids generated in this study are listed in supplementary materials and methods. The RNA sequencing datasets generated during this study will be made available at NCBI Gene Expression Omnibus. For further information and requests for resources and reagents kindly contact, Roger Foo.

