## Supplementary methods and figures for "The novel conserved NAD^+^-binding micropeptide SGHRT regulates mitochondrial function and metabolism in human cardiomyocytes"

Vinh Dang Do *et al*.

**This PDF file includes:**

Materials and Methods

Figures S1-S6

Captions for Tables S1-S6

Captions for Movies S1-S4

References

**Other Supporting Online Material for this manuscript includes the following:**

Tables S1-S6 (.xlsx)
Movies S1-S4

### Materials and Methods

#### Generation of *MYH6*-GFP reporter line

EGFP cassette with kanamycin selection was inserted into BACs for *MYH6* (RP11-834J17, BacPac) immediately before the initiating Methionine (ATG) using recombination (Quick & Easy BAC Modification Kit, KD-001, Gene Bridges GmbH). The Tol2 transposon cassette with Ampicilin selection mark was inserted into the loxp site of the BAC in the backbone using recombination. Ten million H1 cells were cultured in medium containing 20% KO serum replacement, 1 mM l-glutamine, 1% non-essential amino acids, 0.1 mM 2-mercaptoethanol and 8 ng $ml^{-1}$ of basic fibroblast growth factor (b-FGF) in DMEM: F12 for 6 days, dissociated into single cells with TrypLE™ Express (Life Technologies), and electroporated with 20 micrograms of Tol2 transposes and 100 micrograms of Tol2/EGFP modified Transposon-BACs. After electroporation, cells were re-suspended in conditioned medium with 10 µM ROCK inhibitor Y276329 (Y27632 (Stemcell Technologies, 72302). ROCK inhibitor was added for the first 48 hours after electroporation. 50 µg/ml geneticin (Gibco, 10131035) was added for selection of positive clones 72 hours post-electroporation. Single colonies were picked into 24 well plates for expansion at 14 days after drug selection. Fluorescent *in situ* hybridyzation (FISH) using non-modified BACs as probes was carried out to validate the incorporation of BAC construct into genome of ES cells (Cytogenetics Services, Genome Institute of Singapore) (Fig. S4L). Karyotyping was performed to confirm a normal chromosome pattern (Fig. S4M).

#### Generation of SGHRT knockout ESC lines using CRISPR/Cas9

Plasmid pMIA3 (Addgene plasmid # 109399) was used for CRISPR/Cas9 mediated KO. Two KO hESC lines were generated using dual single guide RNA (sgRNAs) (Table S6) to remove part of *SGHRT* DNA sequence. Single guide RNA sequence were designed using cloud based software tools for digital DNA sequence editing Benchling and CRISPOR. 10 µM sense and anti-sense oligonucleotides were ordered and annealed to generate the 20 nucleotide spacer that defines the genomic target to be modified (SGHRT 5'end and 3'end). pMIA3 was digested with BsmBI and sgRNA spacers cloned after human U6 promoter with T4 DNA ligase (New England Biolabs) following manufacturer's instruction. Ligated constructs underwent transformation using RapidTrans™ Chemically Competent Cells (Active motif). Plasmid was extracted from bacterial culture, purified and Sanger sequenced to confirm successful cloning. Prior to hESC targeting, cutting efficiency of pMIA3-sgRNAs plasmids were tested on HEK293T using the EGxxFP plasmid (pCAG-EGxxFP was a gift from Masahito Ikawa, Addgene plasmid # 50716) (*1*). Best combination of sgRNAs was then used for final targeting of hESCs. The best pair of guides were cloned into a single pMIA3 plasmid digested with NheI & XbaI, using NEBuilder isothermal assembly (New England Biolabs), according to manufacturer's instructions to make the final pMIA3 dual sgRNA plasmid. Human ESCs were dissociated with Accutase (Stemcell Technologies, 07922) and ~$1.5 \times 10^6$ cells were re-suspended in 100 µl P3 Primary Cell kit solution from Lonza (V4XP-3024) and mixed with 10 µg pMIA3dual sgRNA plasmid. To transfect hESC, nucleofection was performed using program CM-113 on the 4D-Nucleofector System (Lonza). Cells were then plated into Geltrex coated 6-well plate with mTeSR medium (Stemcell Technologies, 85850) containing 5 µM Y-27632. After 2 days in culture, cells were dissociated, FACS sorted for RFP positive cells and collected into a tube containingmTeSR medium (Stemcell Technologies, 85850) with 5 µM Y-27632. 500 to 2000 cells were plated into wells of 6-well plate containing the above media. Single cell clones were monitored and upon sizeable growth, colonies were picked and passaged. Genomic DNA was extracted for genotyping and screened for successful knockout. RT-qPCR was performed to validate the knockout of SG*HRT* transcript.

#### Generation of hESC clones with *ATG*-mutated SGHRT using CRISPR/CAS9 base editing

All restriction enzymes were purchased from NEB, unless stated otherwise. PCR reactions were conducted using Q5® Hot Start High-Fidelity 2X Master Mix (NEB, M0494L). Ligations were conducted using isothermal assembly with NEBuilder® HiFi DNA Assembly Master Mix

(NEB, E2621L). Primers and dsDNA fragments were ordered from IDT. Plasmids, pMIA19 & pMIA20, were built for simultaneous expression of dCas9 base editors and sgRNA from a single construct. Plasmids CMV_AncBE4max_P2A_GFP (Addgene #112100) and CMV_ABEmax_P2A_GFP (Addgene #112101), kind gifts from David Liu (*2*), were digested with MluI restriction enzyme respectively. A gBlock carrying the sequence for human U6 promoter, gRNA cloning site with AarI type II RE sites and chimeric guide RNA scaffold was used as a template to amplify a fragment for isothermal assembly using the primers below:

| | |
|---|---|
| 1sg into ancBE & ABEmax F | gcgatgtacgggccagatatAAAAAAGCACCGACTCGG |
| 1sg into ancBE & ABEmax R | tagtcaataatcaatgtcaacgcgtGAGGGCCTATTTCCCATG |

The digested base editor plasmids were ligated with the gRNA expression fragment. The resulting all-in-one plasmids pMIA19-CBE4 and pMIA20-ABE7 (Fig. S6) were used to clone in the gRNAs for base editing and to target the cells as described in the methods section. Plasmids pMIA19 & pMIA20 will be submitted to Addgene.

Human ESCs were dissociated with Accutase (Stemcell Technologies, 07922) and ~1.5x106 cells were re-suspended in 100 µl P3 Primary Cell kit solution from Lonza (V4XP-3024) and mixed with 10 µg plasmid. To transfect hESC, nucleofection was performed using program CM-113 on the 4D-Nucleofector System (Lonza). Cells were then plated into Geltrex coated 6-well plate with mTeSR medium (Stemcell Technologies, 85850) containing 5 µM Y-27632. After 2 days in culture, cells were dissociated, flow sorted for GFP positive cells and 500 to 2000 cells were plated into wells of 6-well plate containing mTeSR medium (Stemcell Technologies, 85850) with 5 µM Y-27632. Single cell clones were monitored and upon sizeable growth, colonies were manually picked and passaged. Genomic DNA was extracted for genotyping and screened for successful base editing. RT-qPCR and Western Blot were performed to validate if there is any change in the expression of SGHRT transcript.

**Stem cell maintenance and differentiation**

Human ESC cell line H1 was maintained using mTeSR medium (Stemcell Technologies, 85850) on 1:200 Geltrex (Thermo fisher, A1413202) coated tissue culture plates and passaged regularly as cell aggregates every 4-5 days using ReLeSR (Stemcell Technologies, 05872 ) or Accutase (Stemcell Technologies, 07922). Two days prior to starting differentiation, cells were dissociated using Accutase (Stemcell Technologies, 07922) and seeded as single cells in Geltrex-coated 12-well plates at seeding density between 450000-500000 cells. Cardiac differentiation was performed following the published protocol by Lian et al (*3*), with modifications as follows. 8 µM of CHIR99021 (Stemcell Technologies, 72054) was added on day 0 and left for 24 hours followed by medium change. On day 3, 5uM IWP2 (Sigma Aldrich, I0536) was added using 50/50 mix of new fresh medium and conditioned medium collected from each well and left for 48 hours. Culture medium from day 0 until day 7 was RPMI1640 (HyClone, SH30027.01) plus B-27 serum-free supplement without insulin (Gibco, A1895601). From day 7 and onwards RPMI1640 with B-27 serum free supplement with insulin (Gibco, 17504044) was used and changed every 2-3 days.

**Purification of hESC-derived cardiomyocytes**

Metabolic selection of cardiomyocytes using lactate-containing medium was carried out to purify hESC-derived cardiomyocytes after differentiation. Purification medium composed of RPMI1640 without D-glucose (Sigma-Aldrich) supplemented with 213 μg/ml L-ascorbic acid 2-phosphate, 500 μg/ml recombinant human albumin and 5 mM sodium DL-lactate (60% w/w). Purification was performed for a maximum of four days. Cardiomyocytes were then dissociated with TrypLE Express (ThermoFisher) 37 ℃ for 15 minutes and plated onto 0.1% Gelatin-coated wells containing RPMI1640 plus B-27 serum-free supplement (with insulin) supplemented with 20% fetal bovine serum (FBS) and 5 μM ROCK inhibitor (STEMCELL Technologies). Medium was replaced with RPMI1640 plus B-27 serum-free supplement on the next day and maintained thereafter.

**Engineered Heart Tissue (EHT) construction**

EHTs were constructed following the protocol described previously (*4*). hESC-derived cardiomyocytes were added into the reconstitution mixture and mixed thoroughly before casting in 2% agarose moulds placed with silicon racks. EHTs were maintained in culture medium containing DMEM, 10% horse serum, 0.1% insulin, 0.1% aprotinin and 1% Penicillin Streptomycin solution. Medium change was performed every two days. From Day 7 onwards, spontaneous and coherent contraction of the EHT strips can be observed.

**Subcellular fractionation of hESCs**

Biochemical fractionation was adapted from previous publication (*5*). After harvesting, hESCs were washed with phosphate saline buffer (PBS) and resuspended in Buffer A containing 10 mM HEPES [pH 7.9], 10 mM KCl, 1.5 mM $MgCl_2$, 0.34 mM sucrose, 10% glycerol, 1 mM DTT, protease inhibitor, phosphatase inhibitor and topped up with $H_2O$ to 10 mL. Triton X-100, with a final concentration of 0.1%, was then added before the cell suspension was incubated on ice for 10 minutes. Upon centrifugation at 1,300 g for 5 minutes at 4 ℃, nuclei was obtained in the pellet (fraction P1) while the supernatant (fraction S1) was subsequently clarified at 20,000 g for 12 minutes at 4 ℃ to obtain the cytoplasmic extract (fraction S2). The nuclei extract was further fractionated by resuspending in Buffer B containing 3 mM EDTA [pH 8.0], 0.2 mM EGTA [pH 8.0], 1 mM DTT, protease inhibitor, phosphate inhibitor and topped up with $H_2O$ to 10 mL. Cell suspension was subjected to sonication (20% amplitude, 10 seconds x 2) to lyse the nuclei before incubation on ice for 30 minutes. Chromatin (fraction P3) was collected after centrifugation at 1,700 g for 12 minutes at 4 ℃ and nucleoplasm (fraction S3) in the supernatant. Enrichment in subcellular fraction was confirmed by Western blot.

**Lentivirus generation**

The 3rd generation lentivirus system was used. Lentivirus vectors were generated from the backbone pLenti-C-mGFP (Origen). CMV promoter was replaced with EF1α and mGFP replaced with blasticidin (BSD) cassette. gBlock of wiltype and alanine-mutated *SGHRT* ORF was cloned after the EF1α promoter into the lentivirus vector, followed by plasmid transformation and bacterial purification. Virus was generated using HEK293T cells and transfected with 7.5 μg of pMDLg/pRRE, 2.5 μg of pRSV-Rev, 2.5 μg of pMD2.G and 10μg lentivirus vector. Viral supernatant was filtered and mixed with Lenti-Pac. Viral particles were pelleted at 3500 g for 25 min at 4 °C and re-suspended in 200 μl DPBS. On the day before transduction, cells were passaged (1:3 ratio) using ReLeSR and plated into 24-well plates. On day 1, cells were fed with 500 μl pre-warmed mTeSR medium (Stemcell Technologies, 85850) containing 6 μg/ml polybrene and incubated at 37°C for 15 minutes, after which 10 ul of viral particles were added to the culture medium and incubated again for 18-20 h. On day 2, similar to day 1, new fresh media supplemented with 6 μg/ml polybrene was replaced and three times the initial amount (30 ul) of viral particles was added to the culture medium. Between day 3 and 4, media was changed without polybrene. From day 5, blasticidin (BSD) (Thermofisher) was added to the media for selection.

**Mitochondria Enrichment**
Mitochondria is enriched from mammalian heart tissues as described by (*6*). Heart tissue were grounded up with pestle and mortar while frozen with liquid nitrogen, suspended in ice cold solution of 250 mM, 0.1 mM EDTA, 1 mM Tris-HCL (pH 7.4), and homogenised with a motorised Dounce homogeniser. The resulting supernatant is then spun at 600 g for 10 minutes with cooling to 4°C to sediment nucleus, and intact cells, then the supernatant is spun at 8000 g to for 10 minutes with cooling to 4°C to obtain the mitochondria enriched pellet. The pellet is then stored frozen (−20°C) in sucrose buffer.

**SGHRT Peptide Mapping**
For analysis of undigested small proteins, mitochondria were lysed in organic lysis buffer [acetonitrile:hexafluoro-isopropanol:water (70:25:0.56:4.44, by vol) containing 20 mM ammonium formate, pH 3.7] (*7*), sonicated in ice water bath for 15 minutes, and centrifuged at 22,000 g with cooling to 4°C to remove insoluble particulates.

For proteolytic digestion, mitochondria enriched pellet was dissolved in 0.2% ProteaseMAX (Promega, V2071), 50 mM ammonium bicarbonate solution. Samples are not reduced or alkylated, since SGHRT contains no cysteine residue. After diluting ProteaseMax to a final concentration of 0.01%, trypsin (Promega, V5111) or chymotrypsin (Promega, V1061) was added, then incubated at 37°C or 25°C for 3 hours respectively. The digested lysate was then desalted with C18 spin tips (Thermo Scientific, 60109-412), speedvac to dryness, and reconstituted in 30 µl of reconstitution buffer (2% acetonitrile, 0.1% formic acid, in water).

LC-MS data was acquired on an UPLC-QTOF system (Agilent 1290 Infinity II + QTOF 6550B). Chromatographic separation was achieved with a linear gradient using mobile phase A (0.1% formic acid in water) and B (0.1% formic acid in acetonitrile) at a flow rate of 0.40 mL/min on a Agilent AdvanceBio Peptide Map, 2.1 x 250 mm, 2.7 µm column maintained at 60°C. For undigested samples, a Luna Omega 1.6 µm Polar C18 100 Å LC Column 100 x 2.1 mm (Phenonmenex) was used instead. A linear LC gradient was set up with percentage of organic solvent as follows: 2.5% at 0–3 minute, 5% at 5 minutes, 14% at 31 minutes, 22.5% at 63 minutes, 60% at 120 minutes, and 95% between 121 and 123 minutes, with flow rate of 0.4 mL/min. Electrospray positive ionization (ESI+) mode was selected, and MS1 was acquired in the 300–1700 m/z range while auto MS/MS was acquired in 300–1700 m/z range. PEAKS Studio v7.5 (Bioinformatics Solutions Inc.) was used for peptide-spectrum matching at the 1% FDR level against respective species specific SwissProt database, with predicted SGHRT sequences manually added.

**Co-immunoprecipitation**
For co-IP with HEK293 cells co-transfected with expression plasmids encoding Wildtype or mutant SGHRT-FLAG and SGHRT-HA, whole cell lysates were prepared in CoIP buffer (20 mM Tris-HCl (pH 8.0), 1% Triton X-100, 2mM EDTA, 150mM NaCl, 0.1% SDS, and Complete protease and phosphatase inhibitor [Roche, Basel, Switzerland]). Immunoprecipitations were carried out using 70ul of anti-FLAG magnetic beads (Sigma) or 100 µl of Pierce™ Anti-HA Agarose beads (ThermoFisher). Tris/Tricine gel electrophoresis was performed using 16.5% Tris-Tricine gels. Standard western blot procedures were per-formed on input and IP fractions using the following antibodies: mouse anti-FLAG (Sigma, 1:1000), rabbit anti-HA (Cell Signalling, 1:1000)

**Western Blot**
Culture and tissue samples were lysed in cold RIPA buffer with protease inhibitor unless otherwise stated. Equal amounts of samples were loaded into SDS-page gel and transferred to PVDF membrane (0.45 µm, Bio-rad) for blotting. Nitrocellulose membrane (0.20 µm, Bio-rad) was used during transfer for small-sized proteins on 16.5% SDS-page gel. Primary antibody used: rabbit anti-C7ORF73 (Novus Biologicals, 1:1000); mouse anti-FLAG (Sigma,

1:1000); rabbit anti-HA (Cell Signalling, 1:1000), mouse anti-GAPDH (Sigma-Aldrich, 1:10000), mouse anti-LAMIN B1 (Abcam, 1:1000), rabbit anti-SDHA (Cell Signalling, 1:1000).

**Amide Hydrogen-Deuterium Exchange Mass Spectrometry (HDXMS)**
SGHRT micropeptide (1mg) in lyophilized form (Pepmic Co., Ltd) was resuspended in 50 µl of 100% DMSO (Sigma-Aldrich, St. Louis, MO) and subsequently diluted to yield 4 mg/ml stock. Hydrogen-deuterium exchange labeling reactions of SGHRT were initiated by diluting 100 pmol (3 µM) of the micropeptide in pure deuterium oxide (Cambridge Isotope Laboratories, Tewksbury, MA) at ~90% final concentration. After 1 and 10 min labeling time, the reactions were quenched by lowering the $pH_{read}$ ~2.5 using chilled trifluoroacetic acid (Sigma-Aldrich, St. Louis, MO). Next, SGHRT micropeptide (3 µM) was mixed with saturating concentrations of FAD (600 µM), NAD (600 µM) and NADH (600 µM), prior to the hydrogen-deuterium exchange reactions. For reference, undeuterated reactions of SGHRT diluted in water were also performed. Each reaction was carried out in triplicates.

The quenched samples (50 µl) were immediately injected into a HDX manager module and subjected to proteolysis for 3 min using immobilized pepsin (Enzymate, Waters, Milford MA), maintained at 12 °C, with a continuous flow rate of 100 µl/min 0.1% formic acid. The peptides generated by pepsin cleavage were captured on 2.1 × 5 mm C18 VanGuard pre-column, and then resolved via ACQUITY BEH C18 reverse-phase liquid chromatography. Peptides were eluted using a gradient of 8-40% acetonitrile in 0.1% (v/v) formic acid at a flow rate of 40 µl/min pumped by nanoACQUITY UPLC™ binary solvent manager. Ultrapure acetonitrile and LC-MS grade water were obtained from Fisher-Scientific-Merck Millipore (Waltham, MA). Vanguard and C18 columns were maintained at 3°C to minimize back-exchange. Peptides were then analyzed on Synapt G2-Si mass spectrometer (Waters, Manchester, UK) acquired in $HDMS^E$ mode. The mass accuracy was continuously checked with 1 µL/min of 200 fmol/µL of Glu-fibrinopeptide as lockmass reference.

Peptides were identified from undeuterated samples using Protein Lynx Global Server software (Waters Corporation, Milford, MA) using sequence of the synthesized micropeptide as the reference database. These identified peptides were then subjected to following parameters: minimum intensity - 2000, mass error (MHP) - 10 ppm, minimum products per amino acid - 0.1, and maximum peptide length – 25. Peptides which passed these quality criteria were then loaded on DynamX 3.0 software (Waters Corporation, Milford, MA) to calculate deuterium uptake values. In addition, full length micropeptide (47 amino acid residues) not subjected to pepsin digestion was also considered to determine the global conformational changes. Mass spectral envelopes and the retention times were manually checked and confirmed for every peptide and deuterium exchange labelling times and centroid values were measured. All reported values are averages of at least two biological replicates each with three independent deuterium exchange experiments and not corrected for back-exchange.

**Proteomics with TG HEK293T Cells and Mouse Tissue Lysate**
TG HEK293T cells were lyzed with buffer consisting of 20 mM Tris-HCl pH 7.5, 150 mM NaCl, 0.2 mM EDTA, 0.1% Triton-X100 and 20% glycerol containing 1X cOmplete, EDTA-free protease inhibitor cocktail (Roche) and 1X PhosSTOP phosphatase inhibitors (Roche). The lysate was incubated at 4 °C for 30 mins with constant rotating to facilitate lysis. The lysate was then centrifuged at top speed for 15 mins at 4 °C and supernatants were transferred to a clean protein LoBind tube (Eppendorf). Heart left ventricles from 6-week mouse C57Bl/6J were homogenized with a mortar and glass douncer in Bind Buffer. The lysate incubate at 4 °C to facilitate lysis. The lysate obtained were centrifuged at 1000 g for 10 mins twice at 4 °C and the supernatants were transferred to a clean protein LoBind tube. HEK cell lysate was

incubated with Anti-FLAG M2 magnetic beads (SIGMA) that were pre-blocked with 1% blocking-grade blocker (Bio-Rad). Samples were rotated for 4 hours at 4 °C to ensure binding of SGHRT-3XFLAG onto beads. Then the supernatant was removed and the beads were further incubated with mouse heart lysate. The beads were then washed three times with wash Buffer (20 mM Tris-HCl pH 7.5, 150 mM NaCl, 0.2 mM EDTA, 0.1% Triton-X100 and 5% glycerol containing 1X cOmplete, EDTA-free protease inhibitor cocktail and 1X PhosSTOP phosphatase inhibitors) at 4 °C. Beads were submitted to the National University of Singapore Proteomics Core facility (SingMass) for protein identification using on-beads digestion and LC-MS/MS.

**Immunostaining**
Cells were fixed in 3.7% formaldehyde for 10 min at room temperature and stored in DPBS. They were permeabilized in 0.2% Triton X-100 for 10 min followed by a pre-blocking step with 2% BSA for 20 min. Primary antibody incubation was performed in DPBS + 10% goat serum or donkey serum overnight at 4 degree and secondary antibody incubation for 1 hour at room temperature. For nuclear staining, cells were stained with 5 mg/ml DAPI at 1: 2000 dilution for 10 min. Antibodies used were mouse anti-FLAG (Sigma-Aldrich, 1:000); rabbit anti-HA (Cell Signaling, 1:500), rabbit anti-SDHA (Cell Signaling, 1:200), rabbit anti-SDHB (Proteintech, 1:250), rabbit anti-SUCLG1 (Sigma, 1:500), rabbit anti-SUCLG2 (Sigma, 1.2 µg/ml), rabbit anti-SUCLA2 (Proteintech, 1:250), rabbit anti-OGDH (Thermo Scientific, 1:500), rabbit anti-IDH2 (Novus biological, 1:400), anti-SLC25A1 (Sigma, 0.75 µg/ml), mouse anti-OCT4 (Santa Cruz, 1:50), Alexa flour 488 goat anti-mouse (Life Technology, 1:1000), Alexa fluor 594 goat-anti-mouse (Life Technology, 1:1000), Alexa fluor 546 goat-anti-rabbit (Life Technology, 1:1000), Alexa fluor 568 donkey-anti-goat (Life Technologies, 1:1000), Alexa flour 647 goat anti-mouse (Life Technology, 1:1000).

**Proteinase K protection assay**
ES cells that are stably expressed SGHRT-FLAG were washed with ice cold Phosphate Buffered Saline (PBS) 2X. 500 µl of MIM buffer (280mM sucrose + 10mM HEPES at pH 7.2) was added and the cells were scraped and collected. The lysis was incubated on ice for 20 min, vortexed with max speed every 5 min and pelleted down at 600 g for 5 min at 4° C. Supernatant was collected and centrifuged at 12,500 x g for 15 min at 4° C. The resulting pellet was resuspended in MIM buffer and spun again at 12,500 x g for 15 min, while the supernatant was discarded. The isolated mitochondria were suspended in MIM buffer. To 20 µl of mitochondrial suspension, 20 µl of MIM buffer with varying concentrations (0%-2%) of digitonin and fixed concentration of Proteinase-K (5 µg/ml and 1 µg/ml) were added and incubated for 15 min on ice. Samples without Proteinase-K and with 1% Triton X-100 were used as controls. PMSF was added to inactivate Proteinase-K after the incubation period. 4X Laemlli buffer was added and subjected to denaturation at 98°C for 10 min. Samples were then used for western blot analysis.

**RNA isolation and reverse transcription quantitative PCR (RT-qPCR)**
RNA was extracted using Direct-zol™ RNA MiniPrep Kit (Zymo, R2060). Cells were directly lysed using Trizol reagent (Thermo Fisher, 15596026). All experiments were performed following the manufacturer's instructions. RNA (250-1000 ng) was reverse transcribed to cDNA using qScript Flex cDNA Kit (Quantabio, 95049-025) with a combination of random primers (Table S6) and oligo (dT). cDNA (1:10) was mixed with PerfeCTa SYBR Green FastMix, low ROX (Quantabio, 95074-05K) and specific primers on a 384-well plate. Real time qPCR was run using ViiA 7 Real-Time PCR System (Applied biosystems). Average Cq was recorded and ΔΔCq method was used to calculate relative gene expression changes. Expression levels of genes were normalized against two housekeeping genes, *GAPDH* and *PPIA*.

**Bulk RNA seq analysis**

Total RNA was extracted using Direct-zol™ RNA MiniPrep Kit (Zymo, R2060) and taken through library preparation to be subjected to Illumina sequencing. RNA integrity number (RIN) is measured with a Bioanalyzer 2100, using the Agilent RNA 6000 Pico kit (Agilent, 5067-1513), to ensure samples brought down for library preparation have a high RNA quality.

Total RNA sequencing is done with the NEBNext Ultra II Directional RNA Library Prep Kit (NEB, E7760), and involves the use of NEBNext Poly(A) mRNA magnetic beads (NEB, E7490) to retain only mRNA with intact polyA tails from the total RNA samples. The mRNA was fragmented using a chemical mix, followed by first- and second-strand cDNA synthesis using random hexamer primers and dUTP mix to allow for strand specificity. “End-repaired” fragments were ligated with a unique illumina adapter. All individually indexed samples were subsequently pooled together and multiplexed for sequencing. Libraries were sequenced using the Illumina Hiseq 4000 sequencing system and paired-end 150 bp reads were generated for analysis.

**iTRAQ-based Quantitative Proteomics**

Cells were rinsed in DPBS, scraped in lysis buffer [8M Urea, 10% glycerol, 1% sodium dodecyl sulfate, 50 mM dithiothreitol, 100 mM Tris-HCL pH 6.8, 1X Protease Inhibitor Cocktail (ThermoFisher Scientific 78430)). Benzonase nuclease (Sigma E1014) was then added to reduce viscosity caused by intact genomic DNA. Protein content was estimated by using a commercial Lowry-based assay (RC DC Protein Assay, Bio-Rad) with bovine γ-globulin (Bio-Rad 5000005) as quantitation standard. For each sample, 100 µg of total protein content was reduced, alkylated and trypsin digested using a gel assisted sample preparation (GASP) (*8*) method. In brief, cell lysates were cast into a 20% polyacrylamide gel plug. The gel plug was then shredded with a spin mesh, washed with 8M urea solution and acetonitrile, then incubated with 50 mM triethylammonium bicarbonate buffer (Sigma T7408) and trypsin (Promega V5111) at 37°C overnight. Tryptic peptides were then extracted from the gel fragments with 5% formic acid in acetonitrile, and then vacuum dried.

After 8-plex iTRAQ (SCIEX, Foster City, CA) labelling, the peptides were desalted with C18 cartridge (Waters Sep-Pak), then fractionated on a Agilent 1290 Infinity I LC system, with XBridge BEH C18 3.5 µm, 3.0 × 150 mm (Waters) reversed-phase column, using mobile reserved-phase A (20 mM ammonium formate in water, pH 10) and mobile reserved-phase B (20 mM ammonium formate in 80% acetonitrile, pH 10). Mobile phase gradient of B was as follows: 10 min of 0% B, 156 min of 0–60% B, hold for 5 min at 86% B, 1 min of 60–100% B, hold for 5 min at 100% B, 1 min of 100–0% B, and hold for 14 min at 0% B. The flow rate was set at 0.5 ml/min, and fractions were collected in 1-minute interval. All 175 fractions were then concatenated (*9*) into 30 pooled fraction for LC-MS.

For LC-MS, all 30 pooled fractions were analysed through a nano-LC (Eksigent nanoLC Ultra and cHiPLC-Nanoflex in Trap-Elute configuration) TripleTOF 5600 system (SCIEX, Foster City, CA) under information dependent mode. MS1 was acquired between 400–1800 m/z in high resolution mode (>30,000), while MS/MS was acquired in high sensitivity mode (resolution >15,000) with “adjust CE when using iTRAQ Reagent” turned on. Peptide search was done on ProteinPilot software version 5.0 with the Paragon algorithm, against the Swissprot protein database for *Homo sapiens* (March 2019). The search was further processed by the Progroup algorithm to group redundant proteins that have the same peptides identified. The user-defined search parameters were as follows: Sample Type—iTRAQ 8plex; Cysteine Alkylation—Acrylamide; Digestion—Trypsin; Special Factors—none; Instrument—TripleTOF 5600; Species—Homo sapiens; Search Effort—Thorough; ID Focus— Biological modifications; FDR Analysis—Yes; Background Correction—No; Bias Correction—Yes;

Modified Data Dictionary or Parameter Translation—Yes. The LC-MS method was described previously (*10*).

**Metabolomics profiling**

hES-CMs were subjected to the targeted metabolomics analysis by IC-MS for determination of TCA cycle intermediates. Cells were rinsed with ice-cold PBS one time. 3 ml of ice-cold PBS was added to plates and cells were scraped off the plate and collected into 15ml polypropylene centrifuge tube. Another 3 ml of ice-cold PBS was added to rinse plates to collect remaining cells into the previous tube. Tubes were centrifuged at 1,000 rpm for 5 min at 4°C. Supernatant was aspirated and 300 µL of ice-cold deionized water containing 0.6% formic acid was added to the cell pellets. Cell pellets were resuspended and 30 µL of lysate was aliquotted for protein measurement. 270 µL acetonitrile was then added for cell lysis, and the mixture was vortexed to ensure homogeneity. For TCA cycle intermediate analysis, 300 µL of homogenate was spiked with deuterated internal standards (Lactate-D3, Succinic-C13, Fumarate-D2, Malate-D3, Citrate-D4, Glucose-6-phosphate-C13, Fructose 1,6-bisphosphate-13C (Cambridge Isotope Laboratories, Andover, MA)). Methanol was added to precipitate protein while the supernatant was dried under nitrogen gas. The dried extract was reconstituted in 200 µL of water and used directly for analysis. The analytes were separated using an AS 11-HC column (Dionex, 0.4x250mm, 4 µm, IonPac) on the IC. The IC run was performed at a flow rate of at 0.02 mL/min with an initial gradient of 5 mM of potassium hydroxide (KOH), then increased to 20 mM in 0.5 min, 25 mM in 7 min, 50 mM in 8.5 min, 70 mM in 11 min and 90 mM in 15.5 min. Suppression technology was enabled so that the IC can couple to the Thermo Q Exactive Plus mass spectrometer (Thermo Fisher Scientific, MA) by converting the KOH to pure water. An additional ionization support spray made from methanol with 0.6% formic acid acted as supplement flow to the mass spectrometer. All compounds were ionized in negative mode using electrospray ionization. Absolute quantitation of both organic acids and glycolytic intermediates were done by comparing the ratios of the metabolites to internal standards, against an external calibration curve. Data acquisition and analysis were performed on an Xcalibur software and Trace Finder 4.1 General Quantitative software.

**Seahorse assay**

OCRs of intact cells were measured using Agilent Seahorse XF Cell Mito Stress Test in an XFe96 Extracellular Flux Analyzer (Agilent). A total of 30,000 hES-derived CMs were seeded on Seahorse XF Cell Culture Microplate coated with 0.1% Gelatin in RPMI and B27 with insulin (Fig. 4, I and J) or in CM maturation media comprising DMEM supplemented with 2% foetal bovine serum, 1x non-essential amino acids and 1x GLutaMAX in the presence or absence of 100µM fatty acid solution containing 100 µM of oleic acid and palmitic acid in a 1:1 ratio (Fig. 4, L and M) at 37°C for 72 h. The cells were then incubated with Seahorse XF Base Medium supplemented with 1 mM pyruvate, 2 mM glutamine, and 10 mM glucose at 37°C for 1 h. Three basal OCR measurements were taken, followed by sequential injections of 2 µM oligomycin, 0.4 µM FCCP and 0.8 µM of Rotenone and Antimycin, taking three measurements following each treatment. For normalization, direct cell count was performed. Basal OCR, ATP-linked respiration and maximal respiration were calculated according to the manufacturer's instructions

**Fatty Acid treatment, ATP quantification and TMRM assay**

4-week and 8-week hES-CMs were cultured in RPMI without glucose and 2% FBS prior to fatty acid treatment. Prior to assessment of ATP and mitochondrial membrane potential (TMRM), cells were maintained for 72h in cardiomyocyte maturation media comprising DMEM supplemented with 2% foetal bovine serum, 1x non-essential amino acids and 1x GLutaMAX in the presence or absence of 100µM fatty acid solution containing 100 µM of oleic acid and palmitic acid in a 1:1 ratio. For ATP assay, adherent cells were lysed in boiling water for 10 minutes with intermittent vortexing. The lysate was spun down at 14000 G for 5 minutes to remove debris and the supernatant was analysed using ATP Determination Kit (ThermoFisher Scientific).

For TMRM assay, cells were seeded in a black clear-bottom 96-well plate and incubated with TMRM (ThermoFisher Scientific) at 250nM for 30 minutes at $37^0$C. Following two washes with HBSS, cells were scanned using Infinite 200 Microplate Reader (Tecan, Männedorf, Switzerland) at excitation 538nm and emission at 574nm wavelengths. For both assays, values were normalised to total protein content.

**Enzymatic assays**

Mitochondria was isolated from 1-2x$10^6$ hES-CMs following steps in above mitochondrial enrichment protocol. The mitochondria was subsequently analysed following instruction in different enzymatic assay kits for Succinate Dehydrogenase (SUCL/SCS) Activity (Sigma, MAK197), Isocitrate Dehydrogenase (IDH) Activity (Sigma, MAK062), α-Ketoglutarate Dehydrogenase (α-KG) Activity (Sigma, MAK189), Pyruvate Dehydrogenase (PDH) Activity (Sigma, MAK183), Succinate Dehydrogenase (SDH) Activity (Sigma, MAK197).

**NAD quantification**

Total NAD (NAD + NADH) was extracted from 1 million of hES-CMs with 500 ul of NADH/NAD extraction buffer by freeze/thawing for 2 cycles of 20 mins on dry ice followed by 10 mins at room temperature. Total NAD was measured according to instructions in NAD/NADH quantification kit (Sigma, MAK037).

**Statistical analyses**

Statistical analysis was performed with Prism Graphpad 8.0. Student's t test was used for comparisons between two groups and one-way ANOVA for comparisons between multiple groups. Quantitative data shown as mean ± s.e.m. Both the label-free and iTRAQ-based quantification results were further analysed with Perseus v1.6.6.0 (Tyanova et al., 2016).

**Table S1. List of SGHRT-interacting proteins identified from IP-MS experiment**

**Table S2. List of differentially expressed proteins in iTRAQ data**

**Table S3. List of differentially expressed genes in RNA-seq of 8-week WT and KO hES-CMs**

**Table S4. List of differentially expressed genes in RNA-seq of D7 WT and KO EHTs**

**Table S5. List of differentially expressed genes in RNA-seq of D18 WT and KO EHTs**

**Table S6. List of oligonucleotides**

**Movie S1. Representative movie of D16 WT EHT in Fig. 4E**

**Movie S2. Representative movie of D16 KO EHT in Fig. 4E**

**Movie S3. Representative movie of D27 WT EHT in Fig. 4E**

**Movie S4. Representative movie of D27 KO EHT in Fig. 4E**

Figure S1

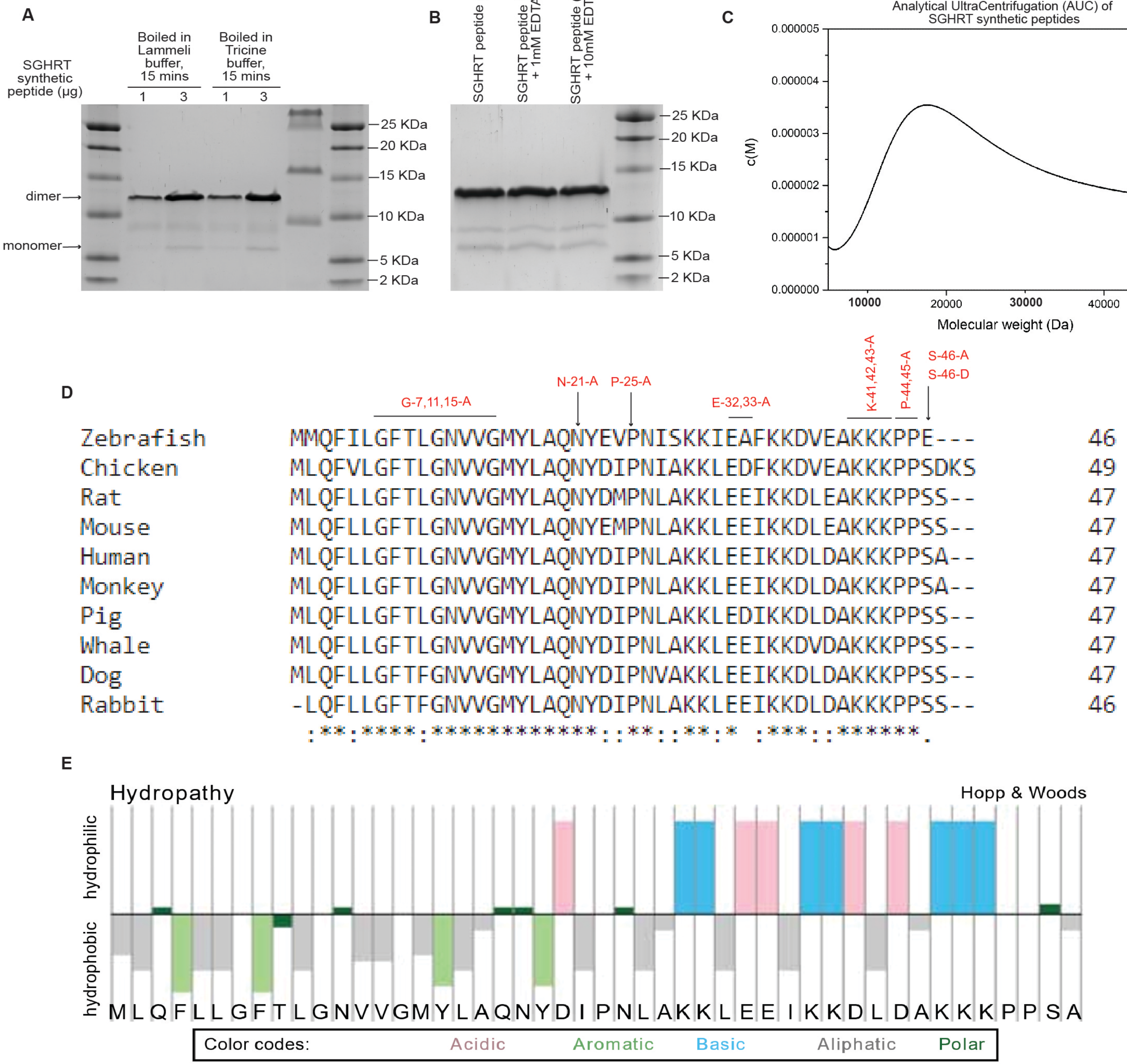
A
SGHRT synthetic peptide (μg)
Boiled in Lammeli buffer, 15 mins
Boiled in Tricine buffer, 15 mins
1
3
1
3
dimer
monomer
25 KDa
20 KDa
15 KDa
10 KDa
5 KDa
2 KDa
B
SGHRT peptide (3 μg)
SGHRT peptide (3 μg) + 1mM EDTA
SGHRT peptide (3 μg) + 10mM EDTA
25 KDa
20 KDa
15 KDa
10 KDa
5 KDa
2 KDa
C
Analytical UltraCentrifugation (AUC) of SGHRT synthetic peptides
c(M)
0.000005
0.000004
0.000003
0.000002
0.000001
0.000000
10000
20000
30000
40000
Molecular weight (Da)
D
G-7,11,15-A
N-21-A
P-25-A
E-32,33-A
K-41,42,43-A
P-44,45-A
S-46-A
S-46-D
Zebrafish MMQFILGFTLGNVVGMYLAQNYEVPNISKKIEAFKKDVEAKKKPPE--- 46
Chicken MLQFVLGFTLGNVVGMYLAQNYDIPNIAKKLEDFKKDVEAKKKPPSDKS 49
Rat MLQFLLGFTLGNVVGMYLAQNYDMPNLAKKLEEIKKDLEAKKKPPSS-- 47
Mouse MLQFLLGFTLGNVVGMYLAQNYEMPNLAKKLEEIKKDLEAKKKPPSS-- 47
Human MLQFLLGFTLGNVVGMYLAQNYDIPNLAKKLEEIKKDLDAKKKPPSA-- 47
Monkey MLQFLLGFTLGNVVGMYLAQNYDIPNLAKKLEEIKKDLDAKKKPPSA-- 47
Pig MLQFLLGFTLGNVVGMYLAQNYDIPNLAKKLEDIKKDLDAKKKPPSS-- 47
Whale MLQFLLGFTLGNVVGMYLAQNYDIPNLAKKLEEIKKDVDAKKKPPSS-- 47
Dog MLQFLLGFTFGNVVGMYLAQNYDIPNVAKKLEEIKKDLDAKKKPPSS-- 47
Rabbit -LQFLLGFTFGNVVGMYLAQNYDIPNLAKKLEEIKKDLDAKKKPPSS-- 46
E
Hydropathy
Hopp & Woods
hydrophilic
hydrophobic
M L Q F L L G F T L G N V V G M Y L A Q N Y D I P N L A K K L E E I K K D L D A K K K P P S A
Color codes: Acidic Aromatic Basic Aliphatic Polar

**Figure S1. Biochemical characterisation of SGHRT micropeptide.**
(**A**) SGHRT synthesised peptides (1µg and 3µg) ran as oligomers in 16.5% Tricine gel.
(**B**) SGHRT synthesised peptides (3µg) ran as oligomers, despite treatment with 1mM and 10 mM EDTA.
(**C**) Analytical Ultracentrifugation (AUC) analysis of SGHRT synthesised peptides suggested that molecular weight of SGHRT peptides exist in the range of 15-20 KDa, with the maximum peak at ~17 KDa.
(**D**) Multiple sequence alignment analysis of mammalian SGHRT micropeptides using Clustal Omega at https://www.ebi.ac.uk/Tools/msa/clustalo/.
(**E**) Hopp and Woods hydropathy plot of STMP1/SGHRT amino acid residues. Generated at https://pepcalc.com/.

Figure S2

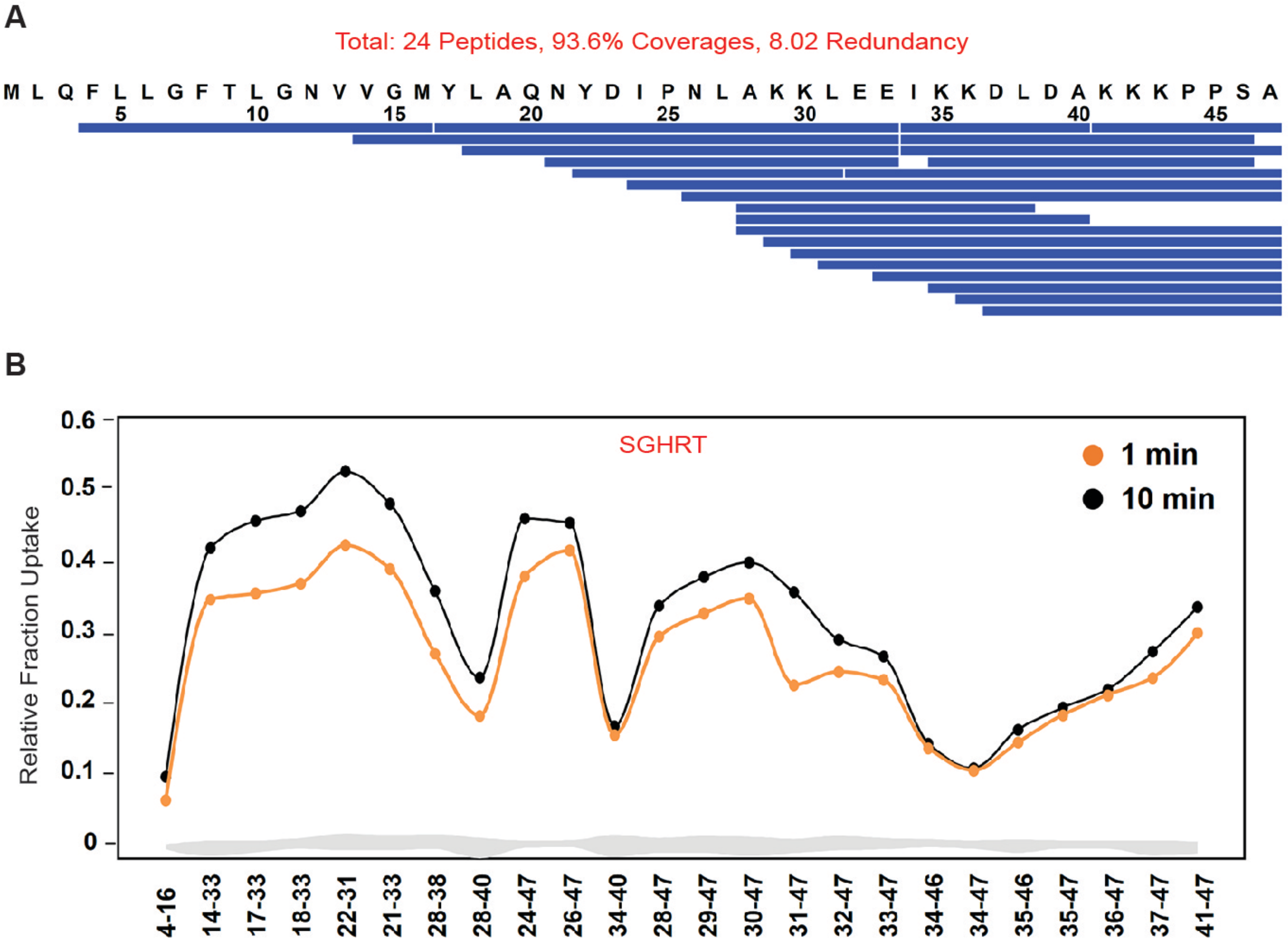
A
Total: 24 Peptides, 93.6% Coverages, 8.02 Redundancy
M L Q F L L G F T L G N V V G M Y L A Q N Y D I P N L A K K L E E I K K D L D A K K K P P S A
5 10 15 20 25 30 35 40 45
B
SGHRT
1 min
10 min
Relative Fraction Uptake
0.6
0.5
0.4
0.3
0.2
0.1
0
4-16
14-33
17-33
18-33
22-31
21-33
28-38
28-40
24-47
26-47
34-40
28-47
29-47
30-47
31-47
32-47
33-47
34-46
34-47
35-46
35-47
36-47
37-47
41-47

**Figure S2. HDX-MS of pepsin-digested SGHRT micropeptides.**
(**A**) Pepsin-digest map of different pepsin-proteolyzed fragments of SGHRT peptides used for HDX-MS experiments.
(**B**) Plot depicting the relative fractional deuterium uptake (Y-axis) of SGHRT for different pepsin-proteolyzed fragments (X-axis) indicated by their residue numbers. Deuterium exchange was performed for 1 and 10 min labelling time scales as indicated.

Figure S3

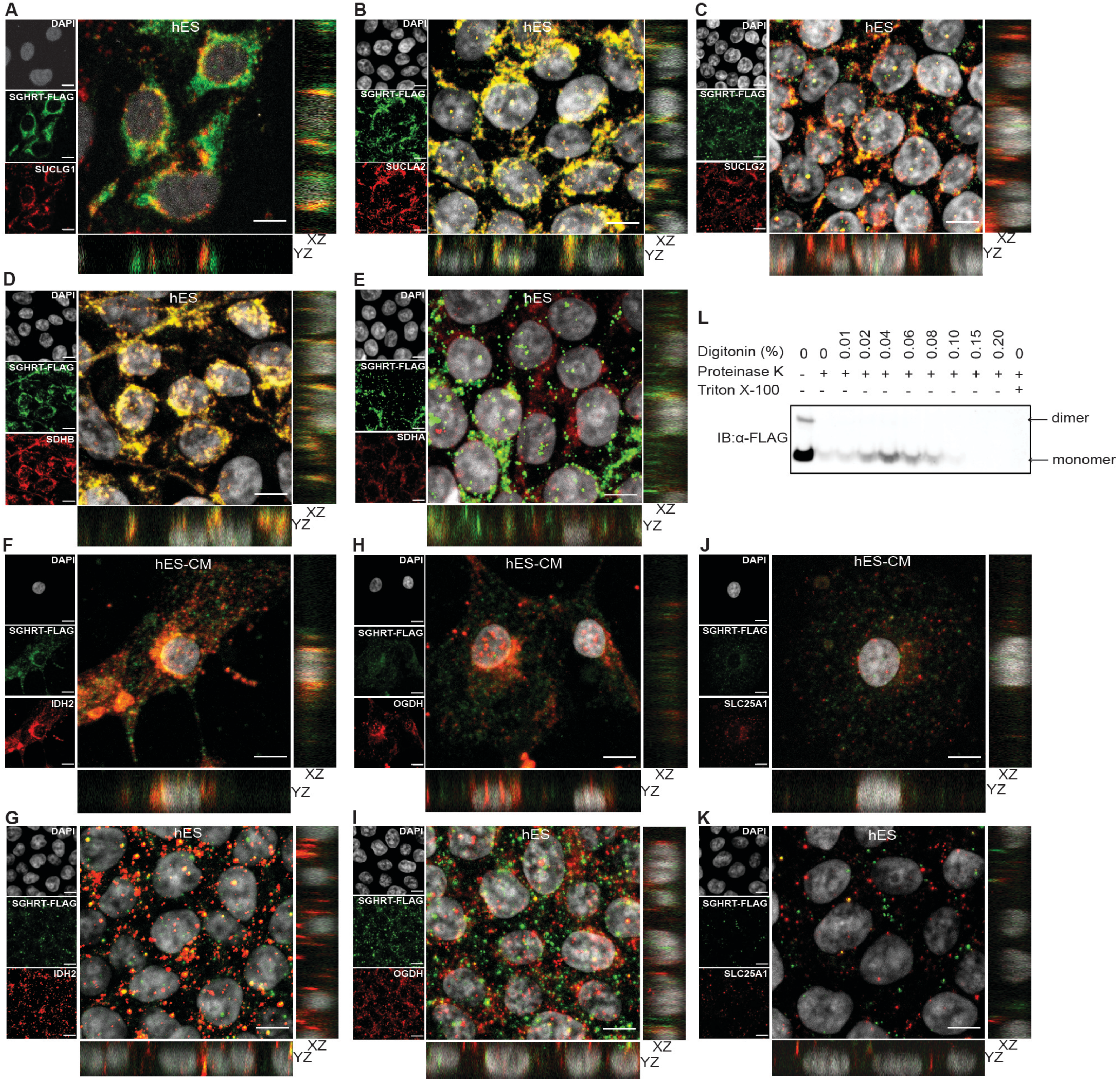
A
DAPI
SGHRT-FLAG
SUCLG1
hES
XZ
YZ
B
SUCLA2
C
SUCLG2
D
SDHB
E
SDHA
F
IDH2
hES-CM
H
OGDH
J
SLC25A1
G
I
K
L
Digitonin (%) 0 0 0.01 0.02 0.04 0.06 0.08 0.10 0.15 0.20 0
Proteinase K - + + + + + + + + + +
Triton X-100 - - - - - - - - - - +
IB:α-FLAG
dimer
monomer

**Figure S3. Characterization of SGHRT-interacting mitochondrial proteins in hES and hES-CMs**

(**A-E**) Immunofluorescence staining of SGHRT-FLAG (green) and TCA cycle proteins (red), SUCLG1 (**A**), SUCLA2 (**B**), SUCLG2 (**C**), SDHB (**D**), SDHA (**E**) in SGHRT-FLAG overexpressed hES cells.

(**F-I**) Immunofluorescence staining of SGHRT-FLAG (green) and NAD+-dependent dehydrogenases (red), IDH2 (**F-G**) and OGDH (**H-I**) in SGHRT-FLAG overexpressed hES-derived cardiomyocytes (**F, H**) and hES cells (**G, I**).

(**J-K**) Immunofluorescence staining of SGHRT-FLAG (green) and NAD+-transporter SLC28A1 (red) in SGHRT-FLAG overexpressed hES-derived cardiomyocytes (**J**) and hES (**K**) cells. Scale bar, 10 μm.

(**L**) Proteinase K protection assay and Western blot of lysates showing SGHRT-FLAG overexpressed in hES cells localised to inner mitochondrial membrane.

Figure S4

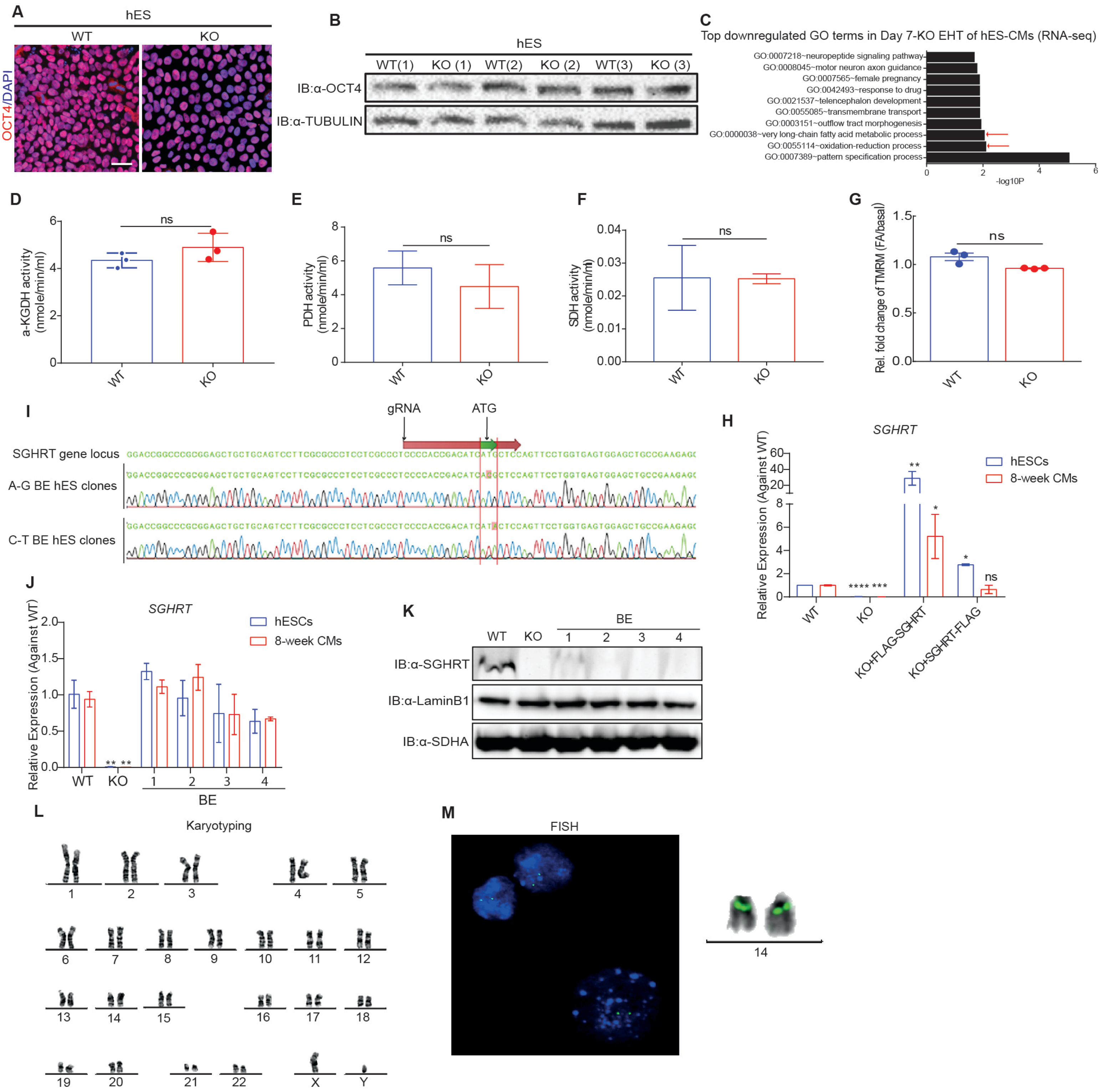
A
hES
WT
KO
OCT4/DAPI
B
hES
WT(1)
KO (1)
WT(2)
KO (2)
WT(3)
KO (3)
IB:α-OCT4
IB:α-TUBULIN
C
Top downregulated GO terms in Day 7-KO EHT of hES-CMs (RNA-seq)
GO:0007218~neuropeptide signaling pathway
GO:0008045~motor neuron axon guidance
GO:0007565~female pregnancy
GO:0042493~response to drug
GO:0021537~telencephalon development
GO:0055085~transmembrane transport
GO:0003151~outflow tract morphogenesis
GO:0000038~very long-chain fatty acid metabolic process
GO:0055114~oxidation-reduction process
GO:0007389~pattern specification process
-log10P
D
a-KGDH activity (nmole/min/ml)
ns
WT
KO
E
PDH activity (nmole/min/ml)
ns
WT
KO
F
SDH activity (nmole/min/ml)
ns
WT
KO
G
Rel. fold change of TMRM (FA/basal)
ns
WT
KO
I
gRNA
ATG
SGHRT gene locus
A-G BE hES clones
C-T BE hES clones
H
SGHRT
Relative Expression (Against WT)
hESCs
8-week CMs
WT
KO
KO+FLAG-SGHRT
KO+SGHRT-FLAG
ns
J
SGHRT
Relative Expression (Against WT)
hESCs
8-week CMs
WT
KO
1
2
3
4
BE
K
WT
KO
BE
1
2
3
4
IB:α-SGHRT
IB:α-LaminB1
IB:α-SDHA
L
Karyotyping
X
Y
M
FISH
14

**Figure S4. Characterization of SGHRT knockout and CRISPR base-edited hES and hES-derived cardiomyocytes**

(**A**) Immunofluorescence staining showing unchanged OCT4 in WT and *SGHRT* KO ES cells. Scale bar, 10 μm.

(**B**) Western blot of total cell lysis of WT and *SGHRT* KO hES showing unchanged OCT4 abundance.

(**C**) GO analysis of RNA-seq data showing top down-regulated terms in D7 KO engineered heart tissues (EHT).

(**D-F**) Unchanged enzymatic activities of alpha-Ketoglutarate Dehydrogenase (α-KGDH) (**D**), Pyruvate Dehydrogenase (PDH) (**E**), Succinate Dehydrogenase (SDH) (**F**), in 4-week WT and *SGHRT* KO hES-derived cardiomyocytes.

(**G**) Quantification of mitochondria membrane potential in WT and *SGHRT* KO hES-derived cardiomyocytes, with and without fatty acid incubation.

(**H**) RT-qPCR result validating expression of *SGHRT* mRNA in WT, *SGHRT* KO, *SGHRT* KO with FLAG-*SGHRT* or *SGHRT*-FLAG rescue, in hES cells and hES-derived cardiomyocytes.

(**I**) Representative Sanger sequencing result confirmed successfully generation of two A-to-G (BE-1 and BE-2) and two C-to-T (BE-3 and BE-4) CRISPR-mediated based edited hES clones.

(**J**) RT-qPCR result showing unchanged *SGHRT* RNA expression in CRISPR BE hES clones.

(**K**) Western blot showing loss of SGHRT in CRISPR BE hES clones.

(**L**) Karyotyping of the *MYH6*-GFP reporter cell line shows a normal karyotype.

(**M**) FISH identified 1 signal incorporated on each allele of chromosome 14 in the *MYH6*-GFP reporter line.

Figure S5

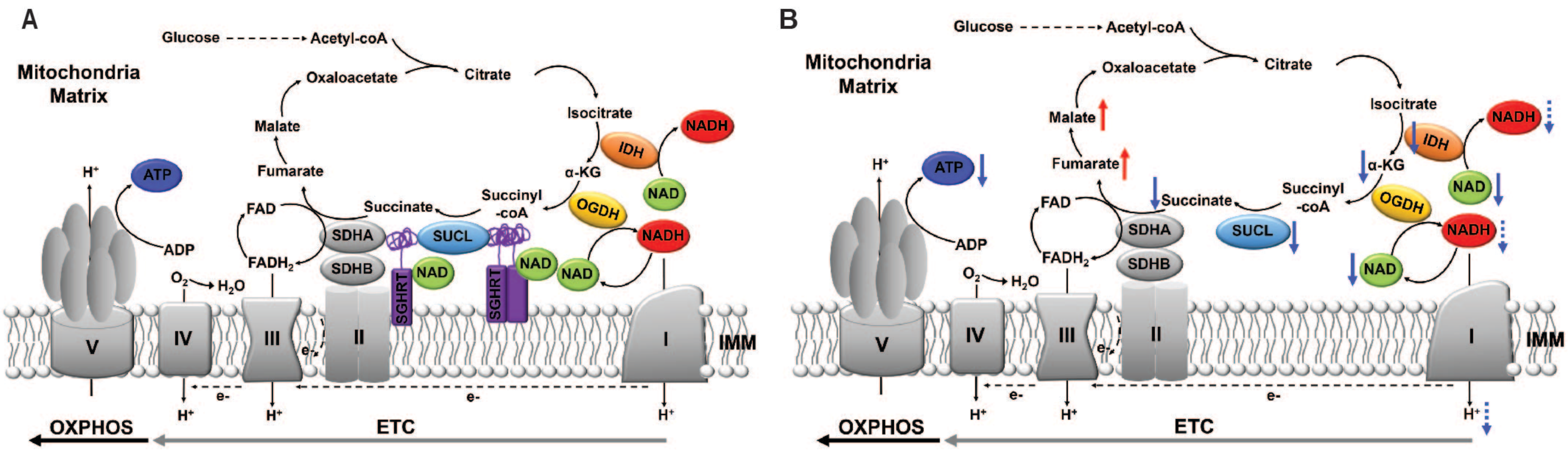

A
Mitochondria Matrix
Glucose
Acetyl-coA
Citrate
Oxaloacetate
Isocitrate
Malate
Fumarate
α-KG
Succinate
Succinyl -coA
IDH
OGDH
NADH
NAD
SDHA
SDHB
SUCL
SGHRT
FAD
$FADH_2$
ATP
ADP
$H^+$
$O_2$
$H_2O$
e-
V
IV
III
II
I
IMM
OXPHOS
ETC
B
Mitochondria Matrix
Glucose
Acetyl-coA
Citrate
Oxaloacetate
Isocitrate
Malate
Fumarate
α-KG
Succinate
Succinyl -coA
IDH
OGDH
NADH
NAD
SDHA
SDHB
SUCL
FAD
$FADH_2$
ATP
ADP
$H^+$
$O_2$
$H_2O$
e-
V
IV
III
II
I
IMM
OXPHOS
ETC

**Figure S5. Graphical illustration of SGHRT micropeptide related signalling events in mitochondria.** (**A**) SGHRT micropeptides (purple) presents as oligomers, and interact with NAD+ (green), and SUCL (blue) and SDH proteins. (**B**) Loss of SGHRT, results in decreased total NAD+, reduction of enzymatic activity of IDH, and reduced α-KG abundance. Decreased α-KG abundance and reduced enzymatic activity of SUCL decreases succinate abundance. Altogether, mitochondrial defects resulting from SGHRT-dependent causes decreased OxPhos and mitochondrial ATP generation.

Figure S6

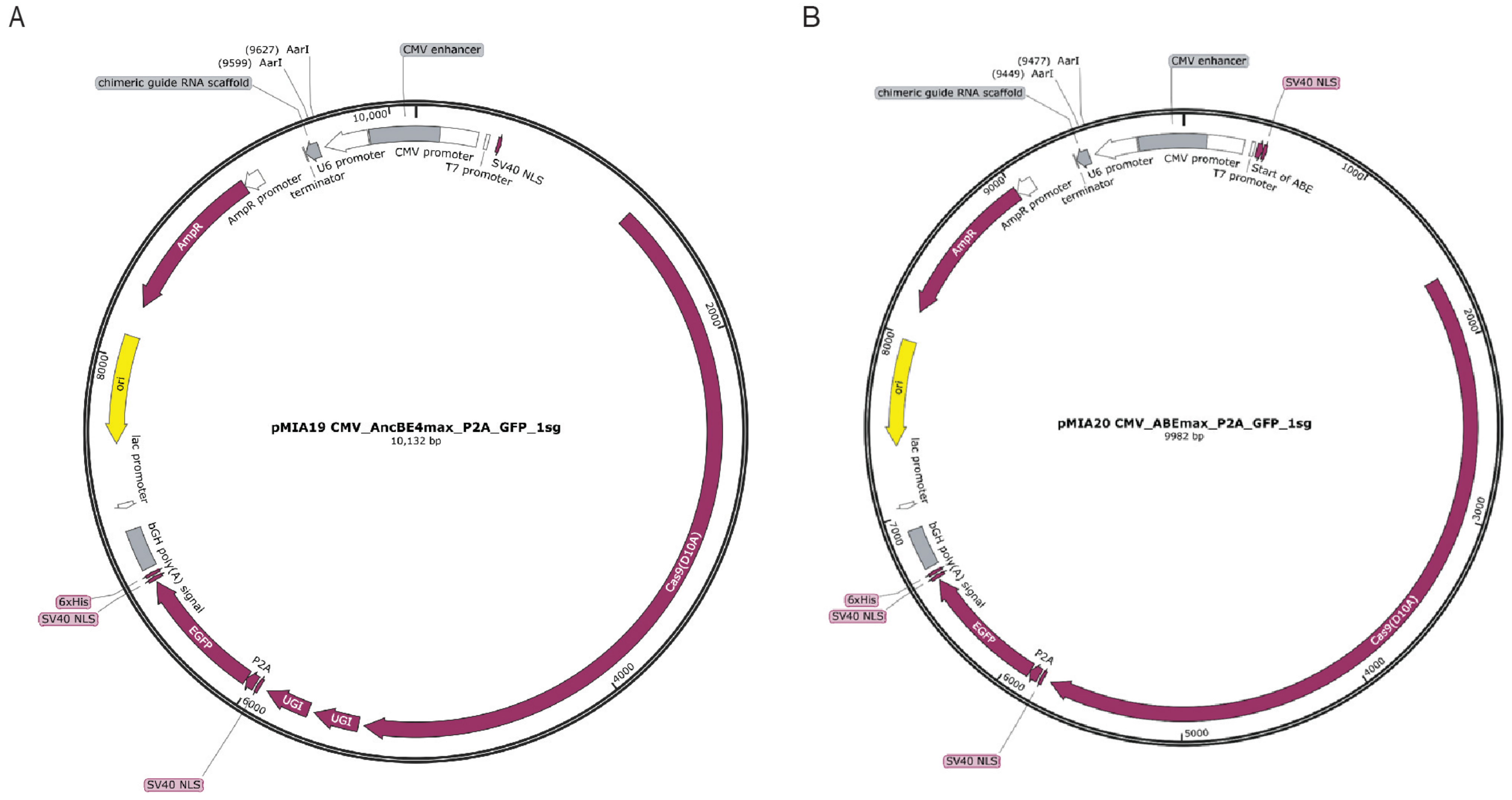
A
(9627) AarI
(9599) AarI
CMV enhancer
chimeric guide RNA scaffold
10,000
U6 promoter
CMV promoter
T7 promoter
SV40 NLS
terminator
AmpR promoter
AmpR
2000
ori
8000
pMIA19 CMV_AncBE4max_P2A_GFP_1sg
10,132 bp
lac promoter
bGH poly(A) signal
Cas9(D10A)
6xHis
SV40 NLS
EGFP
P2A
4000
UGI
UGI
6000
SV40 NLS
B
(9477) AarI
(9449) AarI
CMV enhancer
SV40 NLS
chimeric guide RNA scaffold
U6 promoter
CMV promoter
terminator
T7 promoter
Start of ABE
1000
9000
AmpR promoter
AmpR
2000
8000
ori
pMIA20 CMV_ABEmax_P2A_GFP_1sg
9982 bp
lac promoter
3000
7000
bGH poly(A) signal
6xHis
SV40 NLS
EGFP
P2A
Cas9(D10A)
6000
4000
5000
SV40 NLS

**Figure S6.** Vector maps of all-in-one plasmids pMIA19-CBE4 (**A**) and pMIA20-ABE7 (**B**).
